# The COVID-19 immune landscape is dynamically and reversibly correlated with disease severity

**DOI:** 10.1101/2020.09.18.303420

**Authors:** Hamid Bolouri, Cate Speake, David Skibinski, S. Alice Long, Anne M. Hocking, Daniel J. Campbell, Jessica A. Hamerman, Uma Malhotra, Jane H. Buckner, the BRI COVID-19 Research Team

## Abstract

Despite a rapidly growing body of literature on COVID-19, our understanding of the immune correlates of disease severity, course and outcome remains poor. Using mass cytometry, we assessed the immune landscape in longitudinal whole blood specimens from 59 patients presenting with acute COVID-19, and classified based on maximal disease severity. Hospitalized patients negative for SARS-CoV-2 were used as controls. We found that the immune landscape in COVID-19 forms three dominant clusters, which correlate with disease severity. Longitudinal analysis identified a pattern of productive innate and adaptive immune responses in individuals who have a moderate disease course, whereas those with severe disease have features suggestive of a protracted and dysregulated immune response. Further, we identified coordinate immune alterations accompanying clinical improvement and decline that were also seen in patients who received IL-6 pathway blockade. The hospitalized COVID-19 negative cohort allowed us to identify immune alterations that were shared between severe COVID-19 and other critically ill patients. Collectively, our findings indicate that selection of immune interventions should be based in part on disease presentation and early disease trajectory due to the profound differences in the immune response in those with mild to moderate disease and those with the most severe disease.

## Introduction

The coronavirus-19-disease (COVID-19) pandemic has brought a worldwide focus not only on the severe acute respiratory syndrome coronavirus-2 (SARS-CoV-2), but also on how immunity to this virus both promotes viral clearance and contributes to morbidity and mortality in infected individuals. There is a wide range of disease severity in SARS-CoV-2 infected individuals, ranging from asymptomatic infection to severe COVID-19 requiring mechanical ventilation, and in some cases, to death. Some factors have been identified that are associated with increased disease severity and poor outcome during COVID-19, including age, race, obesity, hypertension, and type 2 diabetes (1-11). However, we still do not understand the biologic factors that contribute to disease severity and outcome. It is becoming clear that not only does the severity of disease vary amongst SARS-CoV-2 infected individuals, but the immune response can also vary widely leading to differing immune landscapes between patients. Therefore, it is important to understand how the immune landscape contributes to COVID-19 severity and outcome. Another important gap in our knowledge is how the immune landscape in COVID-19 resembles or is distinct from that seen in critically ill patients hospitalized for other reasons, since the immune landscape may change in the context of critical illness regardless of its etiology. In particular, it is important to determine if the early immune landscape can be used to inform which COVID-19 patients will have a severe disease course, and would benefit from early interventions.

Although we can learn about immunity to SARS-CoV-2 by assessing a snapshot of the immune response at one point in time, the immune response to infection is dynamic and is best studied over time. Early immune responses to viruses are dominated by the innate immune system, including neutrophils, monocytes, plasmacytoid dendritic cells (pDCs) and natural killer (NK) cells, while adaptive immune responses of T and B cells critical for viral clearance develop over days to weeks. Understanding how these populations change over time and relate to disease trajectory can give insight into the signature of a productive anti-SARS-CoV-2 immune response associated with clinical improvement, and whether immune dysregulation contributes to severe COVID-19. Additionally, early in the pandemic hospitalized patients were treated with a variety of experimental therapeutics, including the antiviral agent remdesivir, cytokine modulating therapies, and plasma from convalescent patients, all with varying efficacy in clinical studies and trials. However, how and if these treatments affect the immune landscape before and after therapeutic exposure has not been described. To address these outstanding and important questions regarding the immune response during COVID-19, we used mass cytometry integrated with detailed clinical data to examine how the immune landscape changes over time in severe and moderate disease through natural progression and recovery, and also in the context of immune intervention.

## Results

### Patient demographics and clinical characteristics

We collected peripheral blood from 59 patients with COVID-19 (52 hospitalized patients and 7 ambulatory outpatients) at the Virginia Mason Medical Center, Seattle, Washington during the months of April and May 2020. Notably, we performed deep longitudinal sampling over the course of disease with an average of 4 time points per subject (Range: 1-18; Figure 1) allowing for detailed immune trajectories of recovery. Patients were classified based on maximum disease severity using a 7-point ordinal scale (OS) representing the following outcomes: 1, not hospitalized with resumption of normal activities; 2, not hospitalized, but unable to resume normal activities; 3, hospitalized, not requiring supplemental oxygen; 4, hospitalized, requiring supplemental oxygen; 5, hospitalized, requiring nasal high-flow oxygen therapy, noninvasive mechanical ventilation, or both; 6, hospitalized, invasive mechanical ventilation; and 7, death (12).Of the hospitalized patients, 24 were classified as having severe disease on the basis of requiring management in a critical care unit (CCU); all required mechanical ventilation (maximal OS≥6), except one who was on high flow oxygen (maximal OS=5). The remaining 28 hospitalized patients were not in the CCU and were classified as having moderate COVID-19, with all requiring supplemental oxygen at some point in their hospital course (maximal OS=3-5). The 7 ambulatory patients had mild disease (OS=2) and did not require hospitalization. For a control group, we also collected blood from 17 hospitalized patients who tested negative for SARS-CoV-2; four of these patients were admitted to the CCU and the remainder to the floor. These patients were age and sex-matched to the hospitalized COVID-19 groups, and were admitted for a variety of conditions including respiratory (n=4), cardiac (n=4), gastrointestinal (n=3), neurologic (n=3) and miscellaneous conditions (n=3).

**Figure 1.**
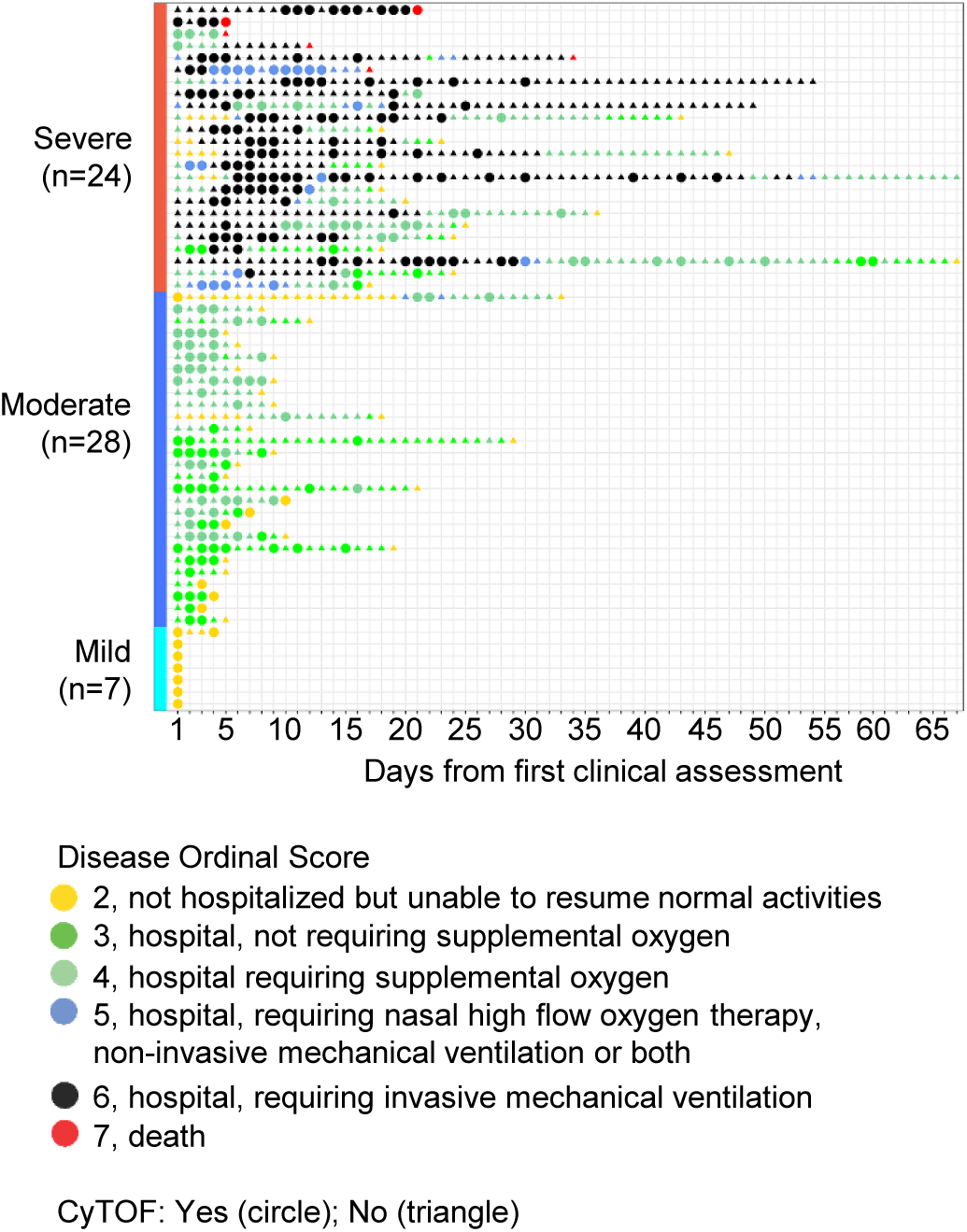
Clinical course and mechanistic data for COVID-19 subjects. Each subject is represented in one row. Subjects are first grouped by severity: severe (red), moderate (blue) and mild (cyan) disease. Subjects are next ranked by highest ever ordinal score (most severe at top), and finally ranked by minimum ordinal score (representing the largest change over time). X-axis: days from first clinical assessment, typically the date of hospital admission. Colored points represent the ordinal score captured daily. No subjects had a score of 1 (recovered) at any point. Dates with CyTOF data available are denoted by circles; dates without CyTOF data are denoted by triangles.

The demographic and clinical characteristics of all the patient groups are summarized in Table 1. There was no significant difference in age or sex composition between severe, moderate and mild COVID-19 groups. Regarding racial distribution, there was an overrepresentation in the severe COVID-19 group of African American (16.7%) and Hispanic (37.5%) individuals based on the Washington state population, which is 78.5% white, 4.4% African American and 13% Hispanic (13). Duration of symptoms at time of presentation was longer in the severe disease group (median 9 days, range 3-22) compared to both the moderate (median 4 days, range 0-27) and mild (median 5 days, range 2-14) groups (p value=0.01). Duration of hospitalization was also significantly longer in the severe disease group (median 19 days, range 4-65) compared to the moderate disease group (median 6 days, range 2-28) (p value<0.01), although discharge was delayed for some patients due to restrictions placed on transfers to skilled nursing facility pending viral clearance from nasopharyngeal swabs.

**Table 1.** Cohort demographics and clinical characteristics.

|  | **Severe**<br>**COVID-19**<br>**(n=24)** | **Moderate**<br>**COVID-19**<br>**(n=28)** | **Mild**<br>**COVID-19**<br>**(n=28)** | **Hospitalized**<br>**COVID-19 negative**<br>**(n=17)** |
| --- | --- | --- | --- | --- |
|  | Median (Range) | Median (Range) | Median (Range) | Median (Range) |
| **Age (yrs)** | 61 (31-89) | 67 (34-96) | 54 (27-76) | 67 (30-97) |
| **Number of days**<br>**hospitalized** | 19 (4-65) | 6 (2-28) | NA | 5 (1-34) |
| **Days from symptom**<br>**onset to admission** | 9 (3-22) | 4 (0-27) | 5 (2-14) | NA |
| **Disease score at**<br>**admission** | 6 (3-6) | 4 (2-4) | 2 (2) | 4 (3-4) |
| **BMI** | 30.1 (18.1-54.5) | 29.0 (17.0-54.1) | 25.5 (22.5-37.2) | 25.7 (21.1-69) |
|  | Number (%) | Number (%) | Number (%) | Number (%) |
| **Outcome: discharged** | 18 (75%) | 28 (100%) | NA | 16 (94.1%) |
| **Outcome: deceased** | 6 (25%) | 0 (0%) | NA | 1 (5.9%) |
| **Female** | 12 (50%) | 14 (50%) | 3 (42.9%) | 10 (58.8%) |
| **Race** |  |  |  |  |
| Asian | 1 (4.2%) | 3 (10.7%) | 3 (42.9%) | 1 (5.9%) |
| African American | 4 (16.7%) | 5 (17.9%) | 1 (14.3%) | 0 (0%) |
| Native Hawaiian/<br>Pacific Islander | 0 (0%) | 1 (3.6%) | 0 (0%) | 0 (0%) |
| Native American | 2 (8.3%) | 0 (0%) | 0 (0%) | 1 (5.9%) |
| White | 6 (25%) | 17 (60.7%) | 2 (28.6%) | 14 (73.7%) |
| Unknown/Other | 11 (45.8%) | 2 (7.1%) | 1 (14.3%) | 1 (5.9%) |
| **Ethnicity:**<br>**Hispanic/Latino** | 9 (37.5%) | 2 (7.1%) | 1 (14.3%) | 0 (0%) |
| **Preexisting**<br>**comorbidities** |  |  |  |  |
| Cancer | 1 (4.2%) | 6 (21.4%) | 0 (0%) | 2 (8.7%) |
| Diabetes | 11 (45.8%) | 8 (28.6%) | 2 (28.6%) | 3 (17.6%) |
| Hypertension | 12 (50%) | 19 (67.9%) | 0 (0%) | 6 (35.3%) |
| **Exposure to**<br>**experimental**<br>**medicine** |  |  |  |  |
| Hydroxychloroquine | 7 (29.2%) | 2 (7.1%) | 0 (0%) | NA |
| Remdesivir | 17 (70.8%) | 11 (39.3%) | 0 (0%) | NA |
| Tocilizumab | 8 (33.3%) | 0 (0%) | 0 (0%) | NA |
| Convalescent plasma | 15 (62.5%) | 4 (14.3%) | 0 (0%) | NA |

Chronic medical conditions such as diabetes, hypertension and cancer were common in the hospitalized COVID-19 cohorts. Diabetes was present in 45.8% of the severe group, 28.6% of the moderate group and 28.6% of the mild group. Hypertension was present in 50% of the severe group and 67.9% of the moderate group but absent in the mild group. Cancer was present in 4.2% of the severe COVID-19 group, 21.4% of the moderate group and absent in the mild group. Obesity was also more prevalent in the hospitalized COVID-19 cohort with a median BMI > 29 in both severe and moderate disease groups compared to a median BMI ∼25 in the mild COVID-19 (p value = 0.08) and the hospitalized SARS-CoV-2 negative groups.

Because this cohort was from the early stage of the pandemic in the USA, hospitalized patients received a variety of experimental treatments, including hydroxychloroquine, remdesivir, tocilizumab and convalescent plasma (Supplemental Figure 1). Notably many patients received more than one type of experimental treatment. In the severe COVID-19 group, 7 patients (29.2%) received hydroxychloroquine, 17 (70.8%) received remdesivir, 8 (33.3%) received tocilizumab and 15 (62.5%) received convalescent plasma. Among the moderately ill, 2 (7.1%) received hydroxychloroquine, 11 (39.3%) received remdesivir, and 4 (14.3%) received convalescent plasma. The mild disease group did not receive any of these COVID-19 therapies.

### Elevated white blood cell counts in COVID-19 are driven by an increase in neutrophils, with correlated depletion of plasmacytoid dendritic cells and basophils

We assessed the immune landscape by combining clinical data with mass cytometry (CyTOF) performed on whole blood samples recovered from the clinical laboratory. The CyTOF panel was designed to assess the composition of the innate and lymphocyte compartments and determine the maturation, lineage and activation status of these cell populations (Supplemental Table 1, Supplemental Figures 2-4). To better understand the impact of disease, we performed correlation analysis on the first sample collected for each patient in the COVID-19 cohort (n=59; Figure 2 and Supplemental Figure 5). The heatmap in Figure 2A shows all significant correlations between clinical data (disease severity ordinal score, age, BMI and CBC) and CyTOF immune cell percentages of the total CD45+ (pan-leucocyte marker) cell compartment, whereas the correlation network in Figure 2B focuses only on correlations among major leukocyte populations identified by CyTOF. We found correlations consistent with the current literature. For example, white blood cell (WBC) counts and neutrophil counts were significantly correlated (Figure 2A), not surprisingly given that neutrophils comprise a large proportion of WBC, and both are elevated in severe COVID-19 (14, 15). Neutrophils in both the CBC and CyTOF datasets also inversely correlated with proportions of lymphocytes and T cells (Figures 2A-B) supporting previous reports that the neutrophil-to-lymphocyte ratio is increased in severe COVID-19 (15-18). In addition, both pDCs and basophils negatively correlated with neutrophils, positively correlated with T cells and positively correlated with each other (Figures 3A-E). Together these findings for pDCs and basophils are consistent with recent studies reporting depletion of these cell types in acute COVID-19 (19, 20). Although our CyTOF panel had limited ability to distinguish T cell lineage, T follicular helper (Tfh) cells were assessed. Notably, unlike other T cell populations the percentage of Tfh cells in the memory CD4+ compartment showed a positive correlation with neutrophils, although this did not reach statistical significant (Figure 3F). Taken together these observations indicate that coordinate and counter-acting changes in neutrophils, lymphocytes, pDCs and basophils drive the immune signature of COVID-19.

**Figure 2.**
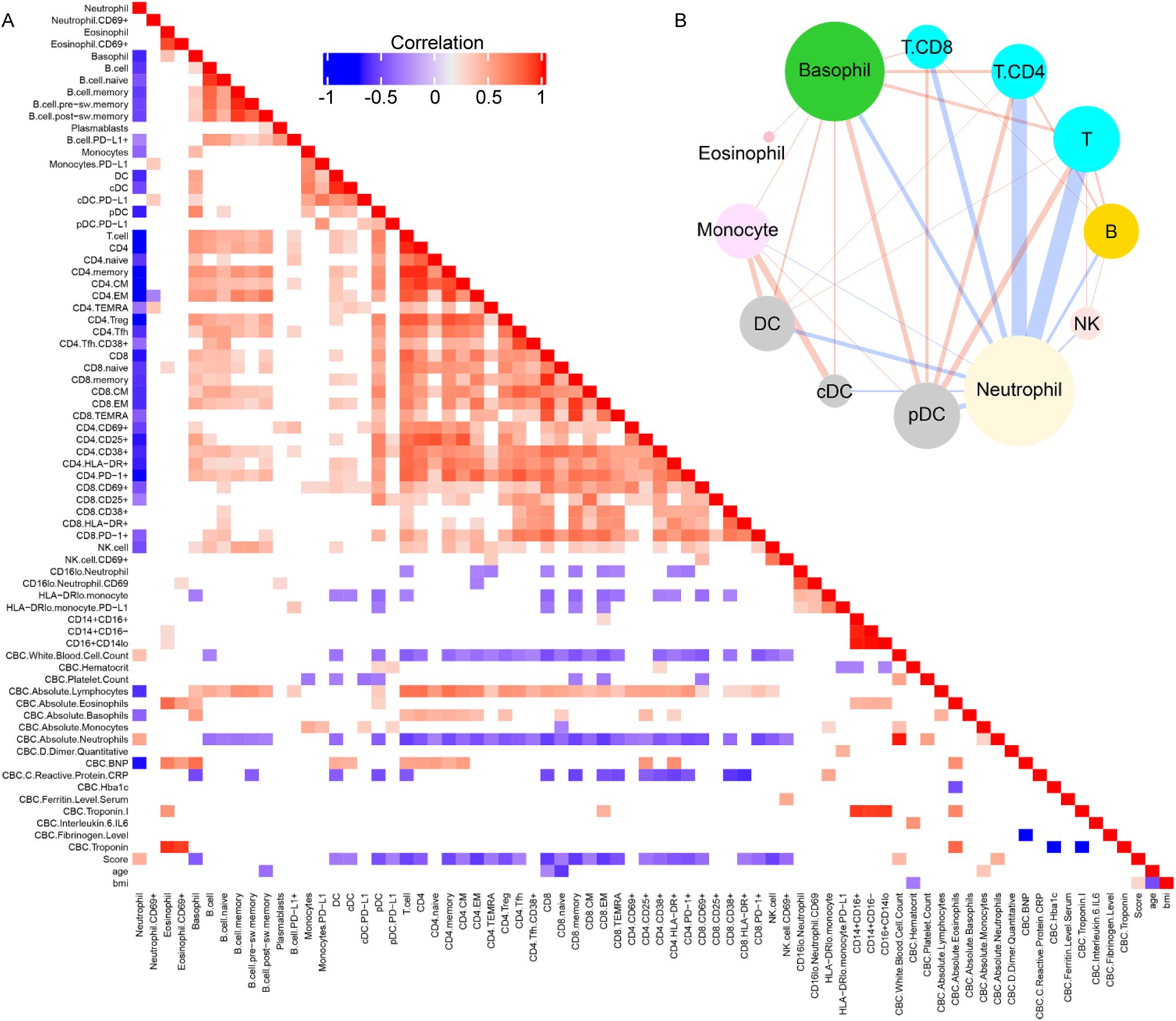
Overview of correlations among cell frequencies and Covid-19 patient characteristics. **(A**) Heatmap visualization of pairwise Pearson correlations with p < 0.05 among ordinal score, age, BMI, CyTOF population frequencies and CBC parameters. Key indicates r value scale for positive (red) and negative (blue) correlations. (**B**) Network map visualization of correlations between CyTOF major immune cell subsets in our mild, moderate and severe COVID-19 cohort. Shown are positive (blue lines) and negative (red lines) Pearson correlations with absolute(r) > 0.35 and p < 0.05. Line thickness corresponds to the strength of association (thicker is stronger). Correlations within major cell populations (same-color nodes) are not shown.

**Figure 3.**
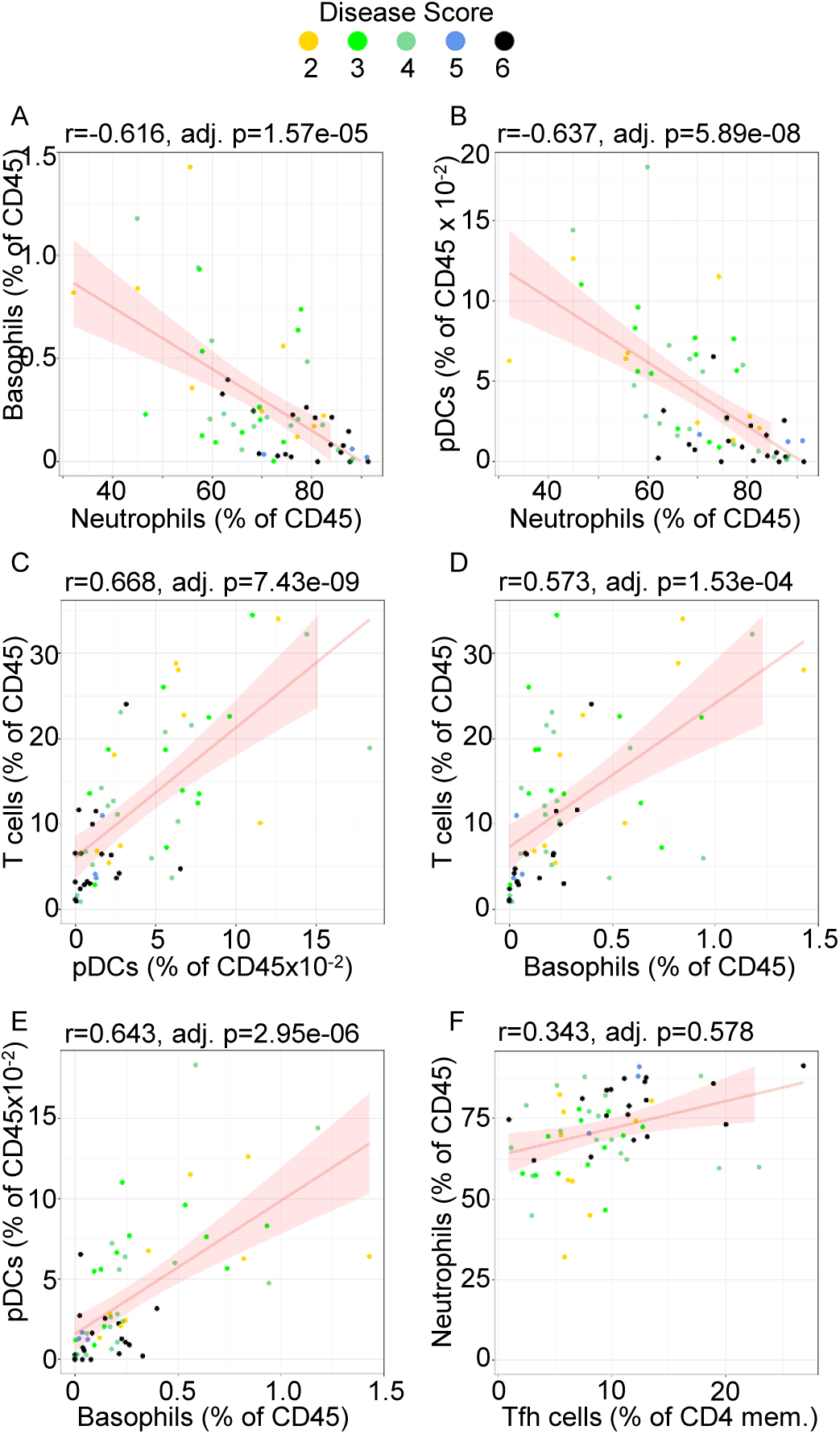
Correlations among immune cell populations in Covid-19 patients suggest disease severity is driven by an increase in neutrophils and a correlated depletion of plasmacytoid dendritic cells (pDC) and basophils. (**A** to **F**) Plots display FDR-adjusted Pearson correlations and linear regression lines with 95% confidence interval shading. Data points are colored according to the ordinal score observed for each patient at admission.

### The immune landscape differentiates individuals based on disease severity

In order to understand whether the immune signature in COVID-19 differed by disease severity we determined the correlation between cell frequency and ordinal score at the time of sampling. Increasing neutrophil frequency was positively correlated with increasing disease severity (Pearson correlation ∼ 0.46, FDR-adjusted p < 0.01), while T cells, NK, pDCs and basophils were lower in severe disease (all FDR-adjusted p-values < 0.005) (Figure 4). To determine if the immune landscape early in disease distinguishes severe from mild disease, we next performed a cross-sectional analysis of our population categorized based on an individual’s highest disease score during the course of their illness using data from the first sample collected for each patient (Figure 5, Supplemental Figure 5). The CBC data showed the greatest difference with disease severity in white blood cell counts with an increase in the absolute neutrophils and monocyte counts and low absolute lymphocyte counts (Figure 5A). However, these CBC results frequently fell within the normal range and notably, the hospitalized COVID-19 negative population showed very similar changes to those seen with severe COVID-19, suggesting that these findings are not unique to COVID-19 but are instead reflective of critical illness.

**Figure 4.**
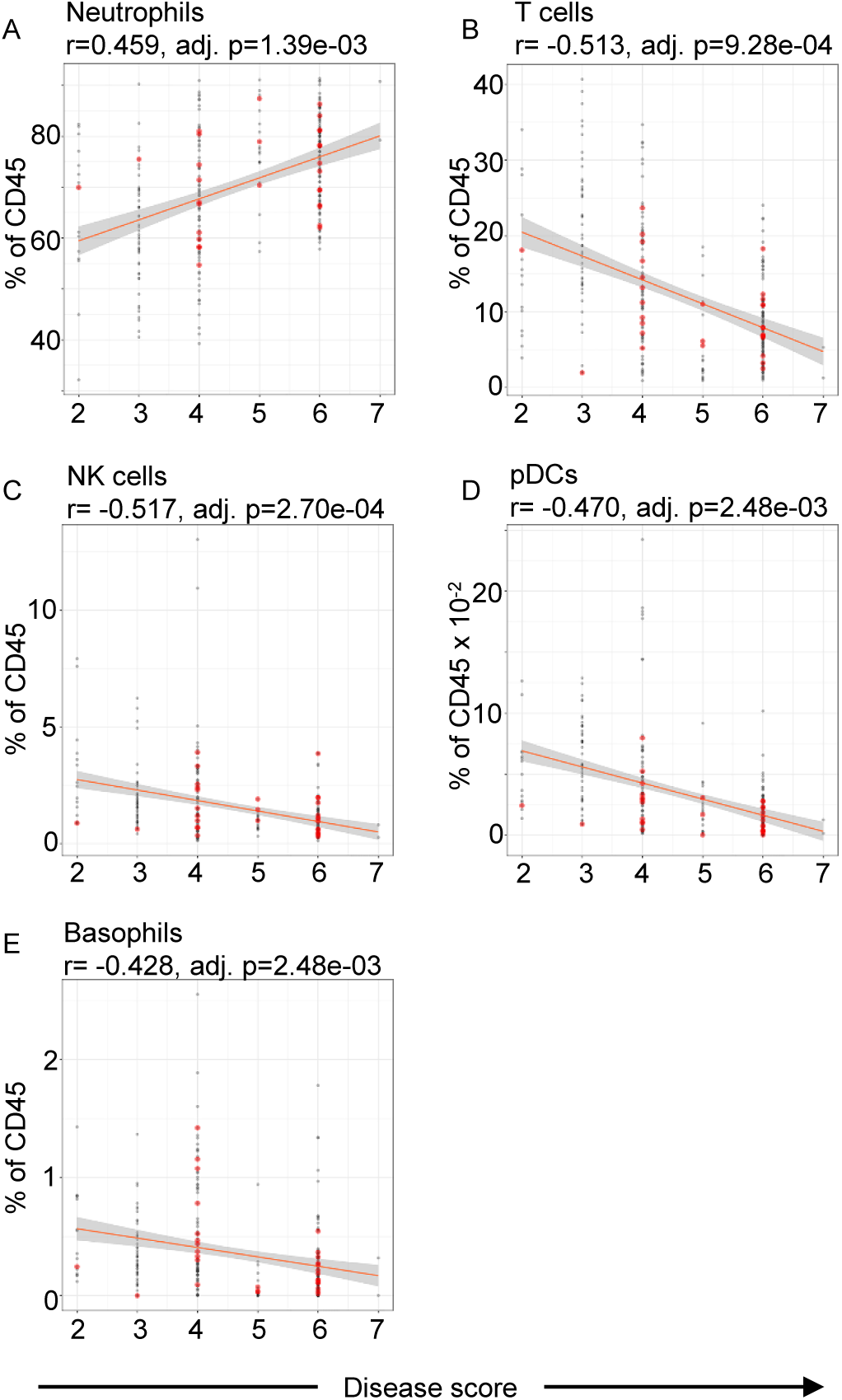
Cross-sectional immune correlates of Covid-19 disease severity. **(A**) In 274 samples from 59 Covid-19 patients, the abundances of neutrophils, T cells, NK cells, plasmacytoid dendritic cells (pDCs), and basophils are highly correlated with disease severity (all p-values FDR adjusted). Red plot points mark values for samples further analyzed in improving versus declining patients (Figure 9).

**Figure 5.**
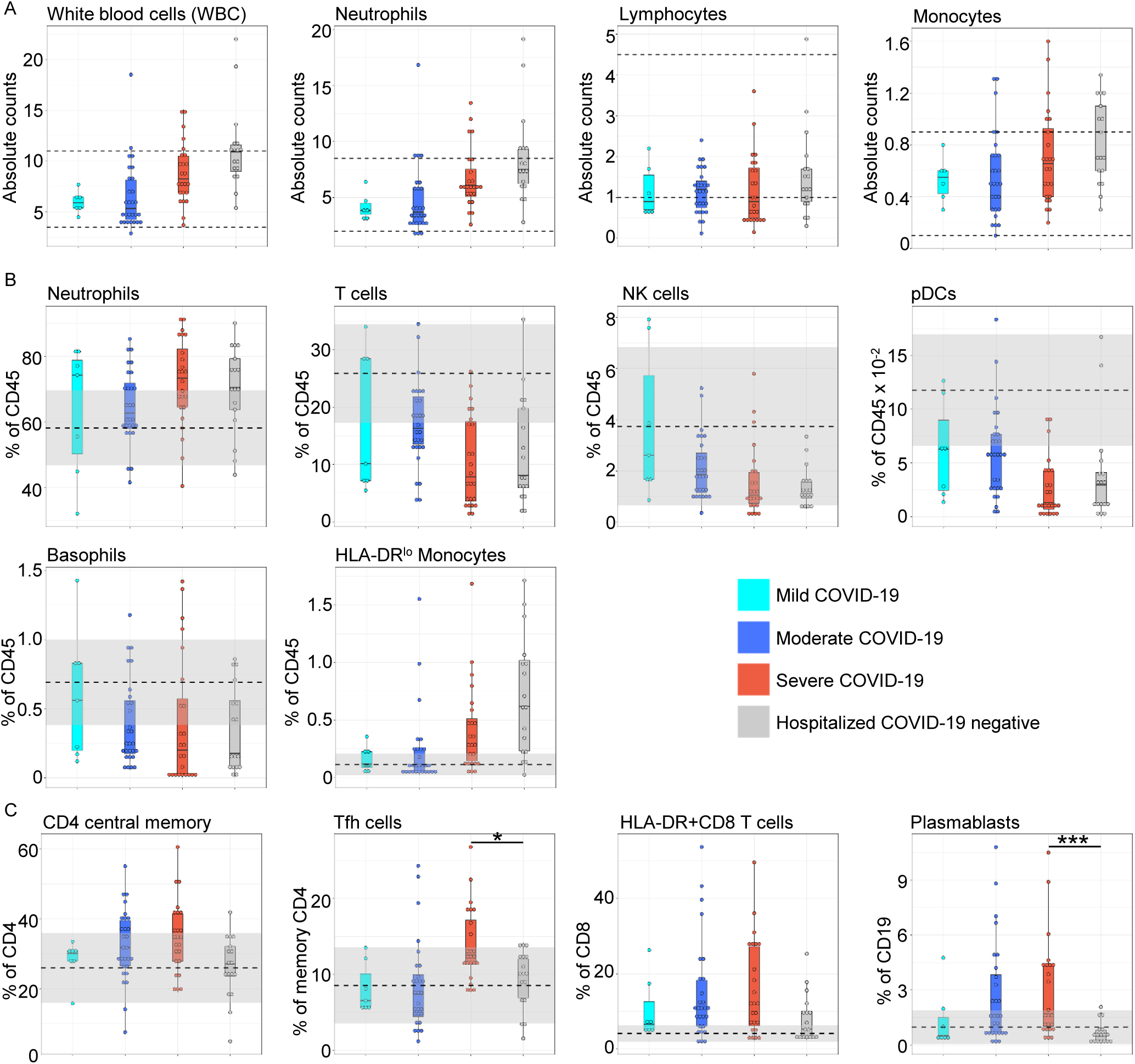
Immune cell frequencies vary by COVID-19 disease severity. **(A)** Clinically-measured CBC absolute count values from day of admission. Dashed black lines mark the clinical laboratories normal ranges. Subjects grouped based on disease severity, mild (cyan), moderate (blue) and severe (red), and SARS-CoV-2 negative hospitalized controls (gray). (**B** and **C**) The relative proportions of immune cell sub-types vary by disease severity. CyTOF cell frequencies based on disease severity expressed as either percentage of all leukocytes (**B**) or percentage of parent population (**C**). Gray bands mark the means (dashed black line) +/-1 standard deviation in 20 healthy control subjects. Asterisks in **C** indicate significance * p<0.05 and *** p < 0.001.

In contrast, the cross-sectional analysis of the CyTOF dataset identified two different patterns of immune alterations in the COVID-19 cohort: those that were also present in the hospitalized COVID-19 negative cohort, and those that were unique to severe COVID-19. Immune cell populations that were similar between severe COVID-19 and hospitalized COVID-19 negative patients correlated with COVID19 disease score at all time points, as shown in Figure 4. Specifically, there was an increase in neutrophils and HLA-DR^lo^ monocytes with a decrease in T cells, NK, basophils and pDCs in severe disease (Figure 5B). Notably, for each of these cell types the changes seen in severe COVID-19 subjects were similar to the hospitalized COVID-19-negative cohort, suggesting that these changes are features of critical illness and not unique to severe COVID-19. Immune alterations unique to severe COVID-19 in this cross-sectional analysis included increases in CD38+ CD8 T cells (FDR-adjusted p = 0.02), Tfh cells (FDR-adjusted p = 0.03) and plasmablasts (FDR-adjusted p = 0.00007) (Figure 5C, Supplemental Figure 5). There were also increases in CD4 central memory T cells and HLA DR+ CD8 T cells although these were not statistical significant after adjusting for multiple testing (Figure 5C).

Unsupervised hierarchical clustering of the CyTOF data for each subject’s initial sample identified three major clusters of patients (Figure 6): a T cell predominant cluster with a relative decrease in neutrophils (cluster A), a cluster with mixed features including a predominance of monocyte, DC and NK cells (cluster B), and a third cluster (cluster C) whose patients had high levels of neutrophils and a relative paucity of other cell types. These clusters generally differentiated individuals based on their disease severity, with more moderate disease courses and good outcome associated with clusters A and B, while those with the most severe disease and death were associated with cluster C. These findings indicate that there is not one single immune signature in COVID-19, but that the immune response differs in individuals based on the ultimate disease severity.

**Figure 6.**
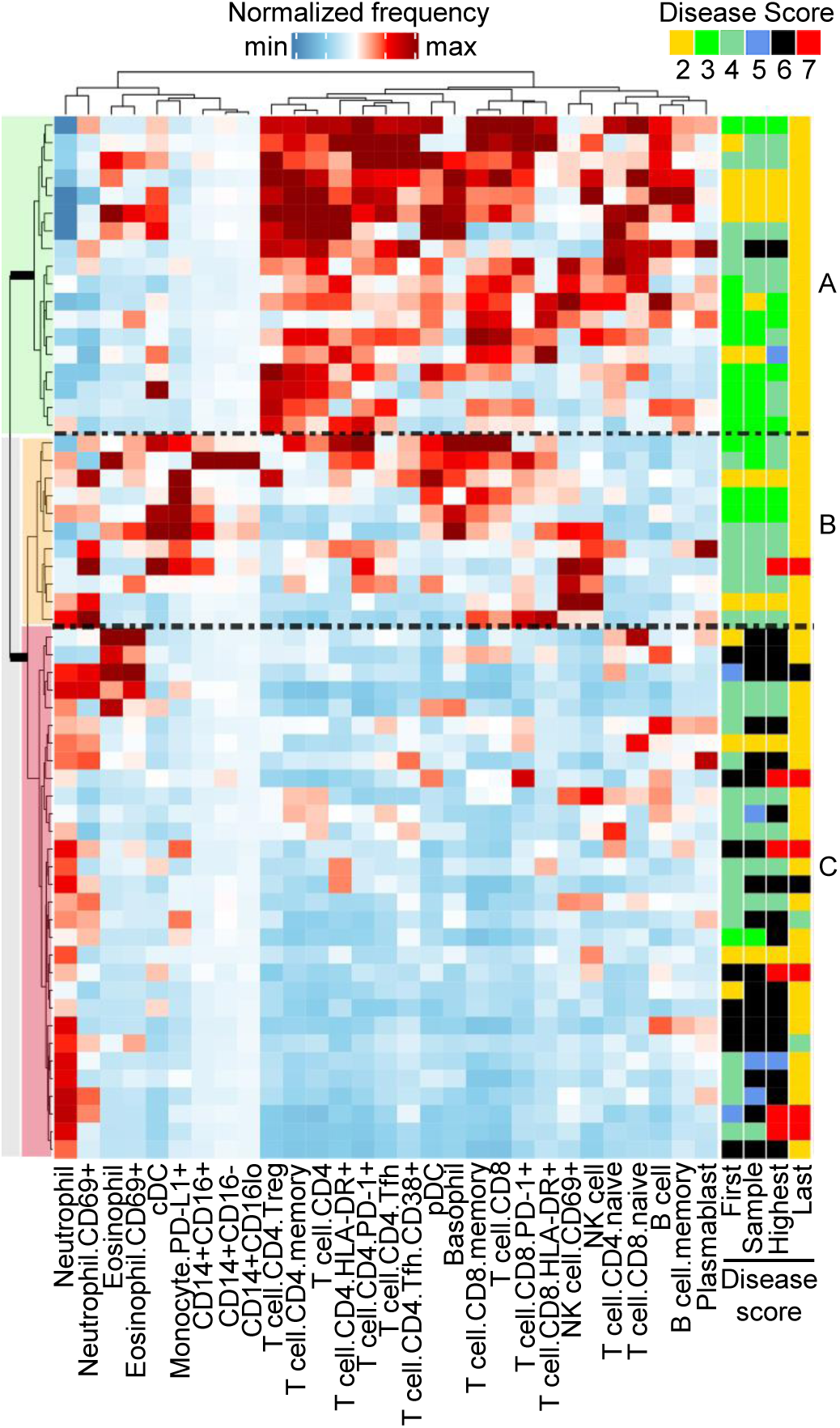
Admission-day sample CyTOF cell frequencies fall into three distinct clusters. Heatmap showing row-normalized Z-scores thresholded at +/-2 (see color key). Disease-severity scores are shown on the right hand-side of the heatmap for the day of admission, day of sampling, maximum score and score at discharge (Disease score key shown at top of heatmap) Clusters are marked A-C at right, and indicated by green, orange, and red highlighting on the dendrogram at left.

### Immune trajectories discriminate moderate and severe COVID-19

To better understand the kinetics and coordinated changes in immune signatures, we tracked immune cell types in the blood over time based on date of admittance to the hospital. We focused on exploring differences in longitudinal analysis of moderate and severe patients based on distinct clustering between these groups as shown in Figure 6. The time course was limited to 15 days post admittance for sufficient and comparable sampling in both the moderate and severe cohorts (Supplemental Figure 6).

Key to understanding features that distinguish moderate from severe COVID-19 is an appreciation of the evolution of the immune response over time, as shown in a UMAP visualization of immune changes with disease severity within cell types of an individual patient (Figure 7). Using gated data from Figures 2 and 3, we focused on specific cell types and markers of innate and adaptive immunity. Figures 5B-D show Loess-smoothed trajectories whereas Supplemental Figures 7-9 show individual and averaged plots. We found that patients with moderate COVID-19 had a dynamic immune response that resolved over time typical of a productive anti-viral response whereas patients with severe COVID-19 had an aberrant immune response, diverging early from that seen in moderate COVID-19 subjects and continuing to diverge beyond the first fifteen days of hospitalization. Specifically in the moderate COVID-19 cohort, there was an early reduction in circulating neutrophils with a concomitant increase in circulating monocytes, total DCs and basophils, with maximal change at 4-5 days post hospitalization (Figure 8A, Supplemental Figures 7-9). In addition, NK cell increases followed these early myeloid cell changes, peaking at 5-6 days post hospitalization (Figure 8A, Supplemental Figures 8-9). In contrast in the severe COVID-19 cohort, these innate cell populations were less dynamic with little variation during the first fifteen days of hospitalization (Figure 8A, Supplemental Figures 8-9). However, it should be noted that this was not the case for all innate cells examined. For example, HLA-DR^lo^ monocytes, which we and others found to be increased in severe COVID-19 (Figure 5B) (21) and are known to be increased in severe inflammatory syndromes such as sepsis (22-24), were more dynamic in the severe COVID-19 cohort than the moderate COVID-19 cohort. HLA-DR^lo^ monocytes in severe COVID-19 subjects increased with time peaking at 5-6 days of hospitalization and then resolved to levels similar to those seen in patients with moderate COVID-19 by day 15 post hospitalization (Figure 8A, Supplemental Figures 8-9). Thus overall, patients with moderate COVID-19 showed a signature of a productive innate immune response in their blood, peaking early after hospitalization, whereas patients with severe disease showed a blunted and delayed innate response.

**Figure 7.**
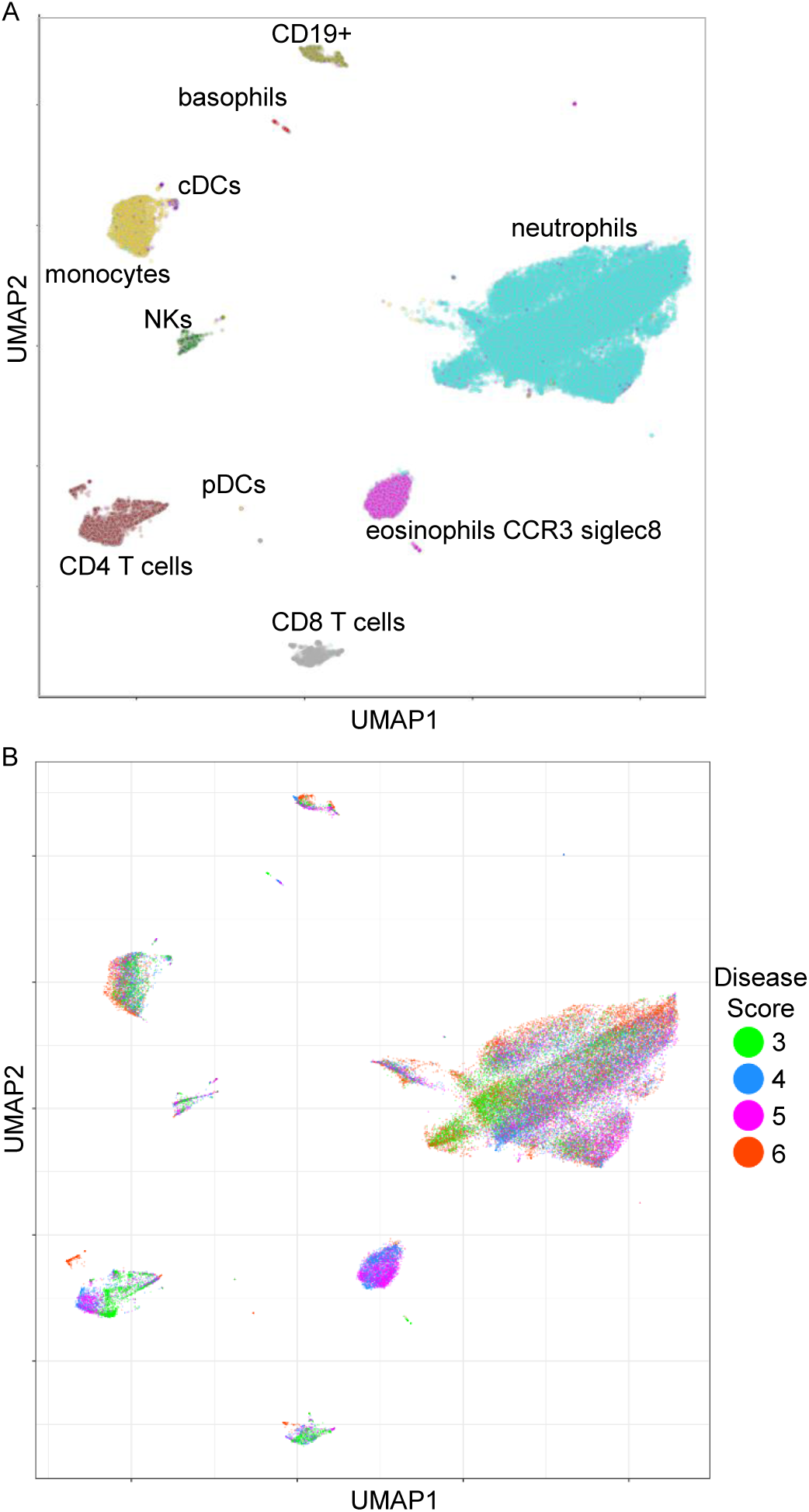
The COVID-19 immune landscape changes over recovery time. UMAP projections of batch-corrected CyTOF probe intensities for four samples from a single COVID-19 patient recovering from a disease severityordinal score of 6 to a score of 3 over a period of 6 weeks (see methods for details).

**Figure 8.**
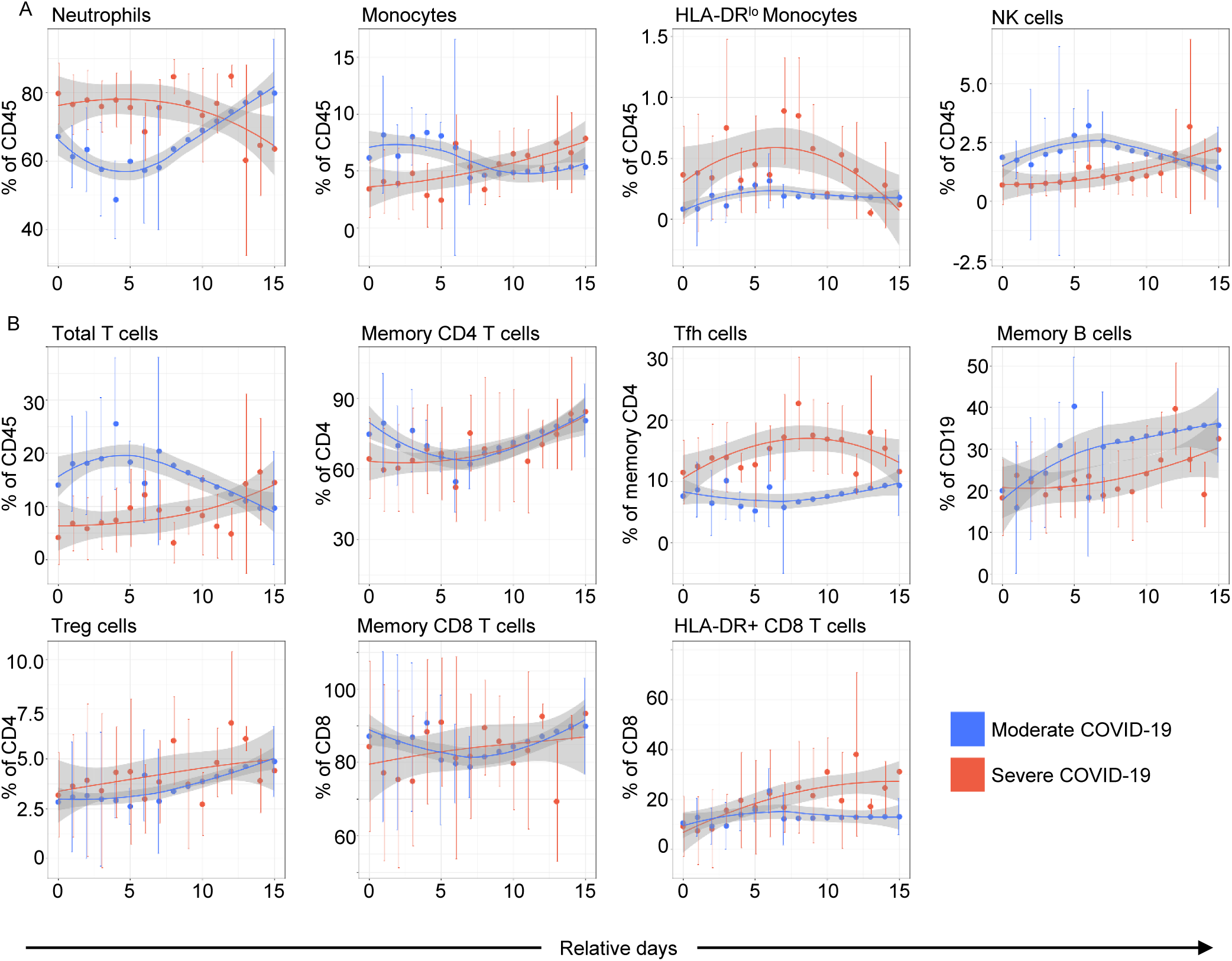
Immune profiles of moderate and severe patients diverge over time reflecting different disease trajectories. Longitudinal plots of gated populations for (**A**) innate and (**B**) adaptive cell types. Days (relative) from first hospitalization are shown. Loess trajectory smoothing was performed on the median values (colored disks) for each group at each time point. Vertical bars indicate +/- 1 standard deviation around the median at each time point. Plot points without vertical error bars are from single data points, or interpolated values used for smoothing.

Adaptive responses also differed between moderate and severe COVID-19 over time (Figures 8B, Supplemental Figures 8-9). In patients with moderate COVID-19, total T cells expanded and contracted consistent with an expected anti-viral T cell response, with a later enrichment of memory CD4 T cells (Figure 8B). Memory B cells increased more robustly over time in the moderate COVID-19 cohort throughout hospitalization, suggestive of sustained interaction with memory CD4 T cells and antibody production. In contrast, patients with severe COVID-19 consistently had lower levels of both T cells and memory B cells over the course of hospitalization suggesting a diminished or delayed adaptive immune response to the virus. The Tfh response in the severe COVID-19 cohort was greater than that of moderate COVID-19 cohort at all time points perhaps indicating unresolved T cell help or Tfh sustained by high IL-6 in critically ill patients. In addition, in the severe COVID-19 cohort, Treg cells as a percentage of total CD4 T cells were increased over time as compared to the moderate COVID-19 cohort (Figure 8B), likely in response to ongoing inflammation due to viral persistence. Consistent with this idea, the percentage of CD8 T cells expressing HLA-DR, a marker of activation, also increased over time in the severe COVID-19 cohort (Figure 8B), as did CD8 T cells expressing CD38 and PD-1 (Supplemental Figure 7B) while total memory CD8 T cells increases were similar between moderate and severe patients (Figure 8B). Overall, our longitudinal analysis revealed that the immune trajectory differs between moderate and severe patients during the first two weeks after initial hospitalization. Patients with moderate disease showed signatures of a productive anti-viral response that resolved within the 2 weeks of the study time, whereas patients with severe disease showed signs of an aberrant response after hospital admittance that persisted for at least the first two weeks in hospital.

### Immune signatures of clinical improvement in patients with COVID-19

To identify key immune cell populations that are associated with either clinical improvement or decline, we focused our analysis on samples taken from individuals before and after a change in ordinal score, reflective of disease severity. We assessed changes in the absolute abundance of immune cell populations by CBC or in the frequency of immune cell subsets in our CyTOF analyses across these key clinical times. We identified subjects that had samples drawn across a score improvement of ≥ 2, or a score decline of ≥ 1 (Figure 9A). This analysis identified several populations whose abundance or frequency was significantly altered upon changes in ordinal score (Figure 9B). Consistent with the lymphopenia observed in severe COVID-19, we found that absolute lymphocytes decreased with clinical decline whereas an increase in the absolute number of lymphocytes was associated with clinical improvement. The increase in lymphocytes was mediated by a general increase in the frequency of naïve and memory CD4+ and CD8+ T cells as well as NK cells, but not B cells. The frequency of pDCs also increased in subjects around the time of clinical improvement, whereas the frequency of neutrophils decreased in improving patients. Longitudinal analysis of individual subjects further demonstrated that changes in the frequency of neutrophils, T cells, NK cells and pDCs could be observed during recovery from severe COVID-19 (Figure 9C). This analysis demonstrates that the immune landscape is dynamic in COVID-19, and that resolution of key features of severe disease resolve co-incident with improvement in clinical status.

**Figure 9.**
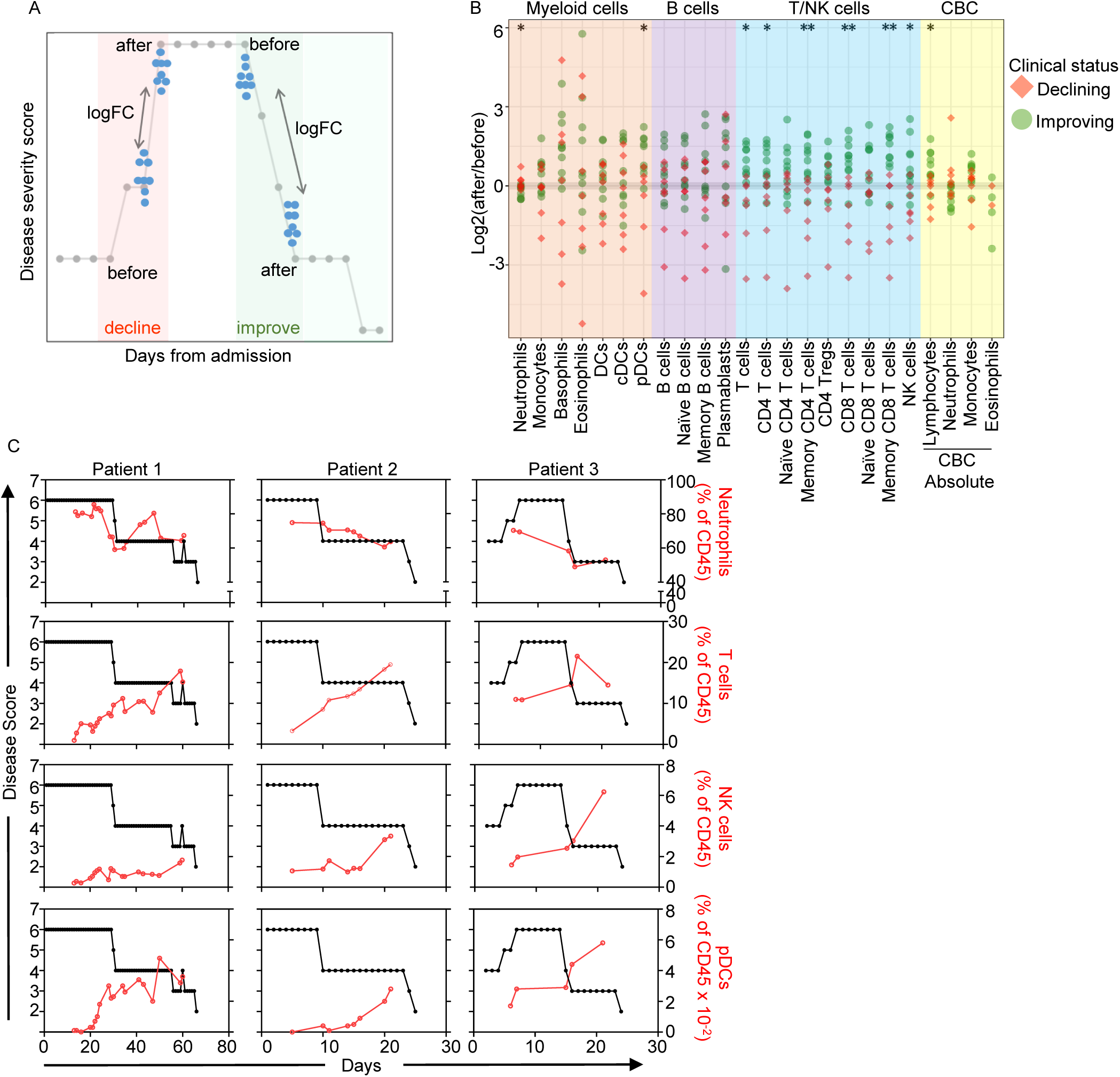
Immune signatures of clinical decline and improvement. (**A**) Schematic outlining approach for identifying immune signatures of clinical decline and improvement focusing on changes in monitored parameters in longitudinal samples taken before and after changes in clinical score. (**B**) Log2-fold change in the indicated cell populations as measured by CyTOF or CBC analysis in longitudinal samples taken before and after improving (green) or declining (red) clinical scores. Asterisks indicates a significant difference in the fold changes (two-tailed, unpaired Wilcoxon Rank Sum FDR-adjusted p < 0.05) between improving (n=7) and declining(n=10) patient groups for the indicated cell populations. (**C**) Longitudinal analyses of the frequency of neutrophils, T cells, NK cells and pDCs vs. clinical score in 3 individual patients.

### Early immune signatures of tocilizumab, but not convalescent plasma, treatment in severe COVID-19 patients

To determine if there were immune signatures of tocilizumab or convalescent plasma treatment, we identified 7 patients treated with tocilizumab and 7 patients treated with convalescent plasma in our cohort who had CyTOF samples both before and after treatment (Supplemental Tables 2 and 3). Notably, these patients all had severe disease and there were stringent criteria for the use of tocilizumab including rapidly escalating oxygen needs combined with an IL-6 level > 20x upper limit of normal (ULN); and CRP >125 mg/dl (ULN, 7). Marked elevations in ferritin, LDH and D-dimer were also weighted in the decision making process. All patients were also treated with remdesivir, with the exception of one patient in the convalescent plasma group. Additionally, 6/7 patients in the tocilizumab group analyzed were treated with convalescent plasma prior to tocilizumab treatment (1-6 days pre tocilizumab). None of the 7 patients in the convalescent plasma group were treated with tocilizumab during the time points analyzed. Also important, patient care was similar between the two groups as the use of tocilizumab in our hospital does not result in alterations to patient care.

We first assessed serum C-reactive protein (CRP) levels in these two groups as a measure of the effectiveness of tocilizumab treatment, which should reduce this marker of systemic inflammation. Indeed, treatment with tocilizumab swiftly reduced serum CRP in all patients (Figure 6A). Serum ferritin was also reduced mainly in those patients with very high concentrations pre-treatment (Supplemental Figure 10A). In contrast, convalescent plasma treatment had no consistent effect on CRP levels (Figure 10A). Therefore, tocilizumab treatment showed an acute clinical signature of reduced inflammation in patients with severe COVID-19, whereas convalescent plasma did not consistently affect these measures.

**Figure 10.**
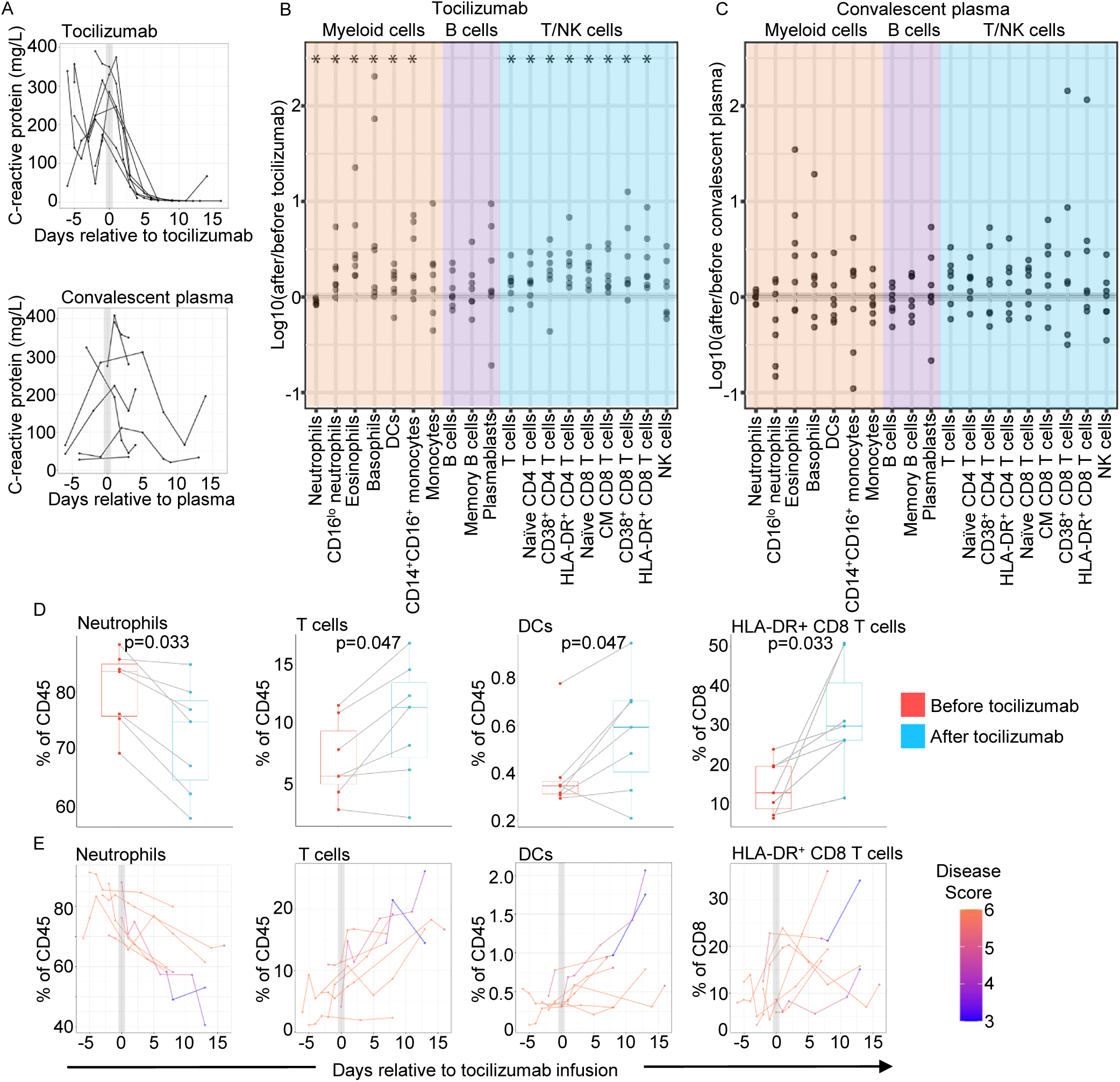
Early immune signatures of tocilizumab but not convalescent plasma in severe COVID-19. (**A**) Serum CRP in patients receiving tocilizumab (top; n=7) or Convalescent plasma (bottom; n=7) measured in clinical labs relative to day of treatment. Each line represents an individual patient. (**B** and **C**) Change in blood immune populations measured by CyTOF after treatment with tocilizumab (**B**) or convalescent plasma (**C**). The fold change in each population for each subject was determined by dividing the percent of each population in the first post- treatment sample at day +2 or more after treatment with the closest pre-treatment sample available as detailed in Supplemental Table 3. All are shown as percent of CD45+ cells unless otherwise indicated. (**D**) Plots showing the percent of the indicated populations in tocilizumab-treated patients before and after treatment, using the time points used for analysis in (**B** and Supplemental Table 3). (**E**) Plots showing all the data points available for tociluzimab-treated patients for the indicated populations shown in (**D** and Supplemental Table 3). Each line represents an individual patient and the color of the line reflects the clinical ordinal score at the time of sampling. *, p<0.05 Wilcoxon matched pairs test, adjusted for multiple comparisons.

We then compared acute changes in immune populations in the blood before and after tocilizumab or convalescent plasma treatment by assessing the closest CyTOF sample before day of treatment (Range: day −4 to day 0) with the first CyTOF sample available after treatment (Range: day 2 to 9 post-treatment). The specific time points used for each individual are shown in Supplemental Table 3. Dividing the post-treatment time point cell frequency by the pre-treatment time point for each patient allowed us to assess the fold change in response to treatment for each patient. In the tocilizumab group there were several populations of immune cells that differed significantly before and after treatment (Figure 10B). In contrast, there were no significant changes after convalescent plasma treatment in the immune cell populations analyzed by CyTOF (Figure 10C, Supplemental Figures 10B-C). The significant changes in response to tocilizumab treatment included a reduction in the percent of neutrophils and an increase in the percent total T cells, eosinophils, basophils, and DCs among CD45+ cells (Figures 10B, 10D, 10E). There were also increases in several CD4 and CD8 T cell subpopulations, and no changes in any B cell populations after tocilizumab (Figure 7B). Moreover, our findings for T cells, B cells, neutrophils and basophils were consistent with our signature of clinical improvement shown in Figure 9. However, there was not complete overlap between the tocilizumab signature and the clinical improvement signature as NK cells and pDCs were not significantly changed by tocilizumab (Figure 10B, data not shown) but were increased in improving patients (Figure 10B). In the tocilizumab group, we also identified increased populations associated with T cell activation, including HLA-DR+ and CD38+ CD4 and CD8 T cells (Figures 10B, 10D, 10E). In summary, we observed a clear acute signature of tocilizumab treatment that shares some but not all features of the immunologic changes seen with clinical improvement, whereas there is no acute change in the immune landscape with convalescent plasma treatment, in patients with severe COVID-19.

## Discussion

A growing literature indicates that the immune landscape is profoundly altered by COVID-19 and differs between individuals dependent on disease severity (18, 20, 25, 26). Whether the immune landscape is a reflection of disease severity, a source of severe disease or a combination of the two is still not fully understood. Here, we utilized recovered samples from the clinical laboratory to rapidly assess peripheral blood cell populations by CyTOF and this data was analyzed in conjunction with clinical laboratories and disease severity scores. Importantly, we were able to collect samples longitudinally among the hospitalized individuals, allowing us to examine the evolution of immune responses through natural progression and recovery, and in the context of immune intervention. Novel aspects of our study included: 1) deep longitudinal sampling allowing for detailed immune trajectories of recovery, 2) a control cohort of moderate and severely ill hospitalized COVID-19 negative patients, and 3) analysis of immune signatures associated with tocilizumab and convalescent plasma treatments.

Notably, we found that at the time of initial sampling the immune landscape in COVID-19 forms three dominant clusters that relate to disease severity. When we examined individual cell populations based on disease severity, we found, as others have, that the neutrophil to lymphocyte ratio is increased in individuals with severe COVID-19 (15-18). Furthermore, this inverse relationship with neutrophils applies to basophils, DC, NK and monocytes, and only modestly with B cells, and is most pronounced among T lymphocytes with the exception of Tfh, which are positively correlated to neutrophil numbers, a finding also consistent with the current literature (18). Interestingly, many changes seen in severe COVID-19 compared to mild and moderate disease were also seen in our hospitalized COVID-19 negative control cohort. Including this unique control group allowed us to identify differences between critically ill patients in general and those infected with SARS-CoV-2. Features shared between the hospitalized severe COVID-19 patients and the COVID-19 negative cohort included increased neutrophils, and decreased T cells, NK cells, pDCs and basophils, and likely reflect active inflammation during critical illness. In contrast, increased Tfh, plasmablasts and evidence of T cell activation were unique to the severe COVID-19 patients and may reflect the anti-viral response in these individuals or unique aspects of the pathology of SARS-CoV-2 infection.

Our longitudinal assessment further allowed us to identify patterns that distinguished severe and moderate disease. Individuals with a moderate disease course showed a pattern consistent with productive innate and adaptive immune response characterized by early and transient increases in monocytes and NK cells with later sustained increases in memory T and B cells. Those with severe disease have features suggestive of a dysregulated immune response characterized by delayed and prolonged increases in Tfh, HLA-DR^lo^ monocytes and activated CD8 T cells. Although the time from first symptom and first sample was delayed in severe patients as compared to moderate, this was not reflected in a simple shift of the immune trajectory. Instead, changes in multiple cell types of moderate subjects were transitory while evidence of persistent activation in different immune cells were progressive and unresolved in severe patients. This suggests that the degree of inflammation or persistence of virus markedly changed the immune landscape over time in severe as compared to moderate disease. Importantly, the persistent features of severe disease are reversed with improvements in clinical score and can be modulated in part with immune interventions, such as IL-6 pathway blockade.

Our findings for the tocilizumab study were intriguing, especially in the light of the recent disappointing results from the first randomized double blind phase 3 trial (27). Our data suggests that changes in the immune landscape after tocilizumab treatment, with the exception of NK and pDC recovery, are consistent with the immune signature of clinical improvement that we identified. Of note, only 4 out of 7 of these tocilizumab-treated patients improved clinically in the acute timeframe of our analysis (within 9 days of treatment) with the remaining 3 patients only showing clinical improvement at later times. This suggests that although tocilizumab treatment induces an acute signature of clinical improvement in both serum CRP and specific immune cell populations, this signature is disconnected from immediate clinical change indicating that the immune changes with tocilizumab may be inadequate to support full recovery. Caveats for our tocilizumab analysis include the small cohort size, and that all but one of these patients were treated with convalescent plasma prior to tocilizumab treatment. Therefore, it is possible that convalescent plasma acts synergistically with tocilizumab to cause the immune signature we identified. Interestingly and in contrast to tocilizumab, we saw no clear immune signature of convalescent plasma within 7 days, suggesting either our cohort was too small to see changes, the immune populations change after the times we analyzed, or convalescent plasma does not act at the level of blood leukocyte populations. It is clear that further investigation is needed to determine if tocilizumab has a therapeutic role in COVID-19, and in what patient population it would be useful and this may be determined in part by the character and trajectory of the immune landscape of the patient.

The demographics of our COVID-19 patients were consistent with published case reports. African Americans and Hispanics were overrepresented in the severe COVID-19 group to the population of Washington State, which is consistent with reports from other states in the USA (10, 11). We also found that type 2 diabetes was more common in those with severe disease compared to moderate or mild disease. Notably, all groups have higher diabetes prevalence than the US or Washington rates (28); the highest prevalence in Washington state is among 65-74 year olds at 21.5%, which is more than doubled in the cohort with severe disease described here.

Diabetes and obesity have consistently been identified risk factors for COVID severity (4-9); reduced T-cell function and chronic inflammation have been postulated as potential mechanisms driving this increased risk (29). In addition, some glucose-lowering agents used in diabetes are known to impact the immune system (reviewed in (30). Full analysis of the differential impact of diabetes and its treatment on our immune signatures is beyond the scope of this work but merits further analysis.

There are limitations to this study. Due to the urgency of the pandemic, we chose to use recovered clinical samples for our study and thus the collection schedule and sample availability was dictated by the treatment needs of the patient. This meant that we did not have the same time points for every patient, and that we could not match between groups the medications that individuals were already taking due to pre-existing comorbidities, some of which may impact the immune responses seen here. In addition, the differences between the mild, moderate and severe COVID-19 groups may reflect the time from disease onset, which significantly varied between these groups, and/or differences in viral burden, which we could not assess. The subjects in our hospitalized control group were not matched to the SARS-CoV-2 positive groups by race, although they are well matched by age.

In summary, we have identified unique features of the immune landscape in moderate versus severe COVID-19 along with features that are common to moderate and severe non-COVID illness. Importantly, our findings indicate that selection of immune interventions should be based in part on disease presentation and early disease trajectory due to the profound differences in the immune response in those with mild to moderate disease and those with the most severe disease. Finally, our characterization of the variety of immune signatures in COVID-19 provides insight into the types of immune interventions that may be beneficial in the treatment of severe disease.

## Methods

### Study design

Using our newly developed 33-parameter CyTOF panel, we characterized the immune response longitudinally in 59 adults with acute COVID-19 including 24 hospitalized patients with severe disease, 28 hospitalized patients with moderate disease and 7 ambulatory patients with mild disease not requiring hospitalization. All COVID-19 subjects were positive for SARS-CoV-2 and our control cohort of 17 hospitalized patients tested negative for SARS-CoV-2. Healthy control subjects were age and sex matched to the hospitalized COVID-19 subjects. Importantly the samples used were collected prior to the start of the COVID-19 pandemic in December 2019. For the hospitalized COVID-19 cohort, longitudinal samples were collected starting as soon as possible after hospital admission, then if feasible, daily for the first week, and then at 3-4 day intervals subsequently (Figure 1). A single sample was obtained at time of first outpatient visit for the ambulatory COVID-19 subjects. A maximum of two samples were obtained from the hospitalized COVID-19 negative control subjects. All assays were run and analyzed in a blinded manner.

### Study approval

Samples from COVID-19 subjects and from hospitalized COVID-19 negative control subjects were recovered from the Virginia Mason Medical Center Central Processing Lab after all tests required for clinical care were complete, under approval by the Benaroya Research Institute (BRI) -approved protocol IRB20-036. All healthy control samples were from healthy subjects in the BRI Immune-Mediated Disease Registry and Repository who had given written informed consent in accordance with the Declaration of Helsinki and according to the BRI Institutional Review Board-approved protocol IRB07109.

### CyTOF staining, acquisition, and subset identification

Peripheral blood was collected from each donor into sterile vacutainer tubes containing the anticoagulant EDTA. Blood cells were washed twice with phosphate-buffered saline (PBS) and stained for viability exclusion with a 100 µM cisplatin solution (Enzo Life Sciences, Farmingdale, New York) for one minute at room temperature. Cisplatin was quenched with five volumes MaxPar Cell Staining Buffer (CSB; Fluidigm, South San Francisco, CA), and the cells then stained with a titered, aliquoted, and frozen cocktail of monoclonal antibodies conjugated to metal isotopes for 20 minutes at 4 °C. Red blood cell lysis was performed using RBC Lysis/ Fixation solution (BioLegend) for five minutes at room temperature followed by a wash with CSB. The resulting leucocytes were fixed overnight at 4°C with MaxPar Fix and Perm Solution (Fluidigm, South San Francisco, CA) containing 125 nM Cell-ID Intercalator-Ir (Fluidigm, South San Francisco, CA). Following fixation, cells were washed with CBS, resuspended in milli-Q water and stored at 4°C until acquisition. All antibodies from Biolegend and BD Biosciences were conjugated to their respective metal isotopes using the Maxpar® X8 Multimetal Labeling Kit (Fluidigm, South San Francisco, CA). Samples were stained within 48 hours of blood draw. Sample stability with the CyTOF assay was established with three COVID-19 samples assayed on the day of collection (baseline, Day 0), one day after collection (Day 1) or two days after collection (Day 2). Populations and markers on populations were standard and based on Staser et.al. (31). All populations and markers with a frequency >5% of live CD45+ cells had a CV<40% between baseline and each time point. This variation was less than that of biological comparisons. MaxPar Four Element Calibration Beads (Fluidigm, South San Francisco, CA) were added to each sample immediately before acquisition. All samples were acquired on a Helios CyTOF3 mass spectrometer (Fluidigm, South San Francisco, CA) with a target cell acquisition of 100,000 live events at a rate of 500 events/second to capture >50 cells per gated population or marker. The CyTOF panel is shown in Supplemental Table 1 and gating strategies are shown in Supplemental Figures 2-4. To determine gates for activation markers such as CD25, CD69 and CD38 on T cells and PD-L1 on myeloid cells, we first analyzed12 samples from 6 subjects with moderate COVID-19 and 6 subjects with severe COVID-19. Gates were set based on a comparison between samples that were clearly highly activated and those that were clearly non-activated. These gates were then applied to all samples in the study and used consistently for all populations analyzed. Specifically, gates for CD25, CD38, CD69, HLA-DR, PD-1 and PD-L1 were the same for all cell types where they were applied. For example, the CD38 gate was the same for CD4 T cells, CD8 T cells and Tfh cells, the CD25 gate was the same for CD4 and CD8 T cells, and the CD69 gate was the same for CD4 T cells, CD8 T cells, NK cells, eosinophils, neutrophils etc. Data was analyzed using a FlowJo software versions 10.6.0 and 10.6.1 (FlowJo LLC, Ashland, OR).

### Data and statistical analysis

#### P-values

Apart from the paired-sample tests in Figure 10, all p-values were calculated using unpaired, two-tailed Wilcoxon Rank Sum tests. In all cases, corrections for multiple testing were performed using the False Discovery Rate (FDR) method. For between group comparisons of the clinical data, p-values were calculated using the Kruskal-Wallis one-way analysis of variance test.

#### Correlation graph

The correlation graph in Fig. 2A was built from the matrix of Pearson correlations in Fig. 2B using the R iGraph package (32)

#### Heatmap

The heatmap in Figure 6 was generated using Euclidean distance and the clustering method Ward.D2.

#### Smoothed time-course graphs

Time-series data from each patient were organized in terms of the relative number of days from the date of the first sample (hereon denoted pseudo-time), and then aligned by first sample. To reduce the potential effects of outlier samples, median values were calculated for each severity category and each day for the samples available. If no samples were available at a given pseudo-time day, we inferred a value using linear interpolation between the before and after pseudo-time points. The vertical bars at each pseudo-time point are equal to one standard deviation from the indicated median value. Plot point with no error bars are those with only one sample or represent an inferred value. Loess smoothing was performed on the median values for each disease severity class using the geom_smooth function in the R ggplot library (33)

#### UMAP

The UMAP plots in Figure 7 were generated directly from the CyTOF signal intensities following archsinh transformation with a co-factor value of 5. To ensure against batch and other potential confounding effects, we specifically selected samples collected and stained in a highly uniform fashion from a single donor and z-score normalized probe intensities for each sample prior to UMAP projection to 2D.

## Author contributions

SAL, CS, UM, JAH, DJC and JHB conceptualized and designed the study. UM and JHB were responsible for subject selection and clinical interpretation. The BRI COVID-19 Research Team obtained IRB approval, collected samples and clinical data, conducted the mass cytometry, and integrated the clinical data with the CyTOF data. SAL, DS and JAH performed the gating for the immune cell populations measured by CyTOF. HB, SAL, DS and JAH performed the computational and statistical analysis. JAH, AMH and JHB wrote the manuscript with assistance from all co-authors. JHB obtained funding and supervised the study.

## Acknowledgements

We would like to acknowledge the Benaroya Family Foundation, the Leonard and Norma Klorfine Foundation, and Glenn and Mary Lynn Mounger for their funding of this project. We also acknowledge the Allen Institute for Immunology for their funding support for the development of the CyTOF panel used in this study. We also would like to thank Carmen Mikacenic for help with obtaining IRB approval, and Henry T. Bahnson for help with statistical analysis. See Supplemental Acknowledgements for consortium details. The graphic abstract was created with BioRender.com.

