## Supplemental Material for "The COVID-19 immune landscape is dynamically and reversibly correlated with disease severity"

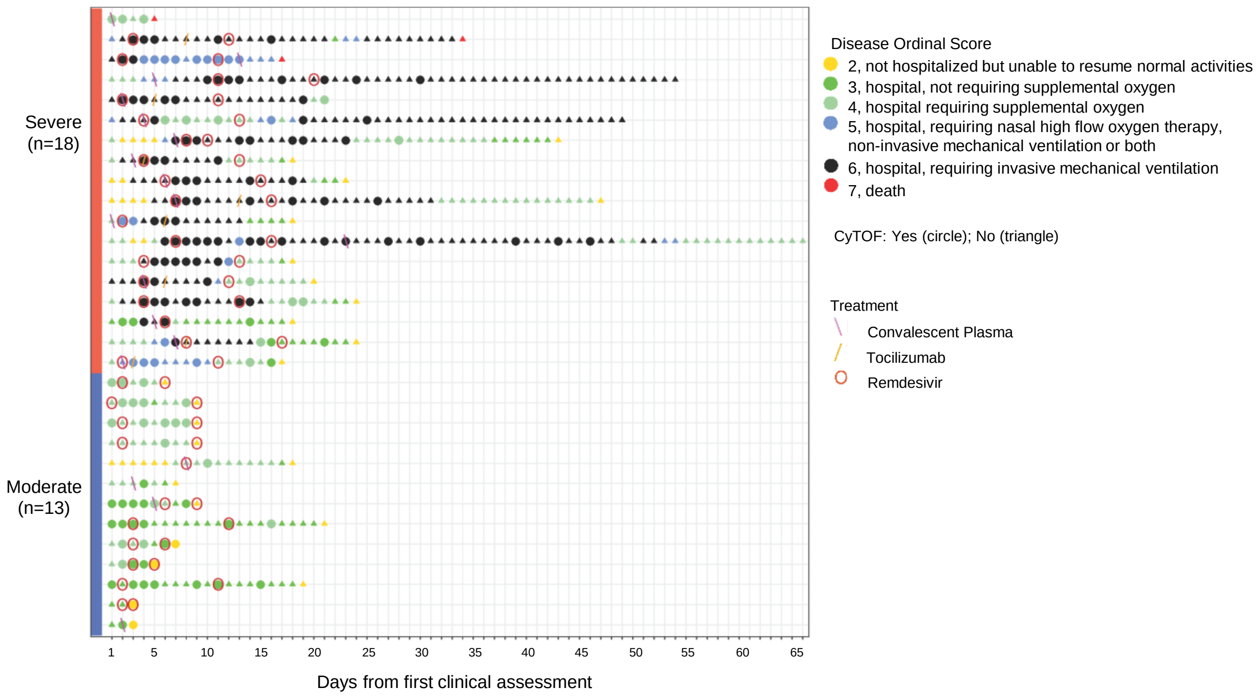

**Supplemental Figure 1. Clinical course, mechanistic data and treatments for COVID-19 subjects.** Subset of COVID-19 cohort that were treated with remdesivir, tocilizumab and/or convalescent plasma (n=13 for moderate COVID-19, and n=18 for severe COVID-19). Each subject is represented in one row. X-axis: days from first clinical assessment, typically the date of hospital admission. Colored points represent the ordinal score captured daily. Dates with CyTOF data available are denoted by circles; dates without CyTOF data are denoted by triangles. Treatments indicated are convalescent plasma (pink backslash), tocilizumab (yellow forward slash) and remdesivir (red open circles).

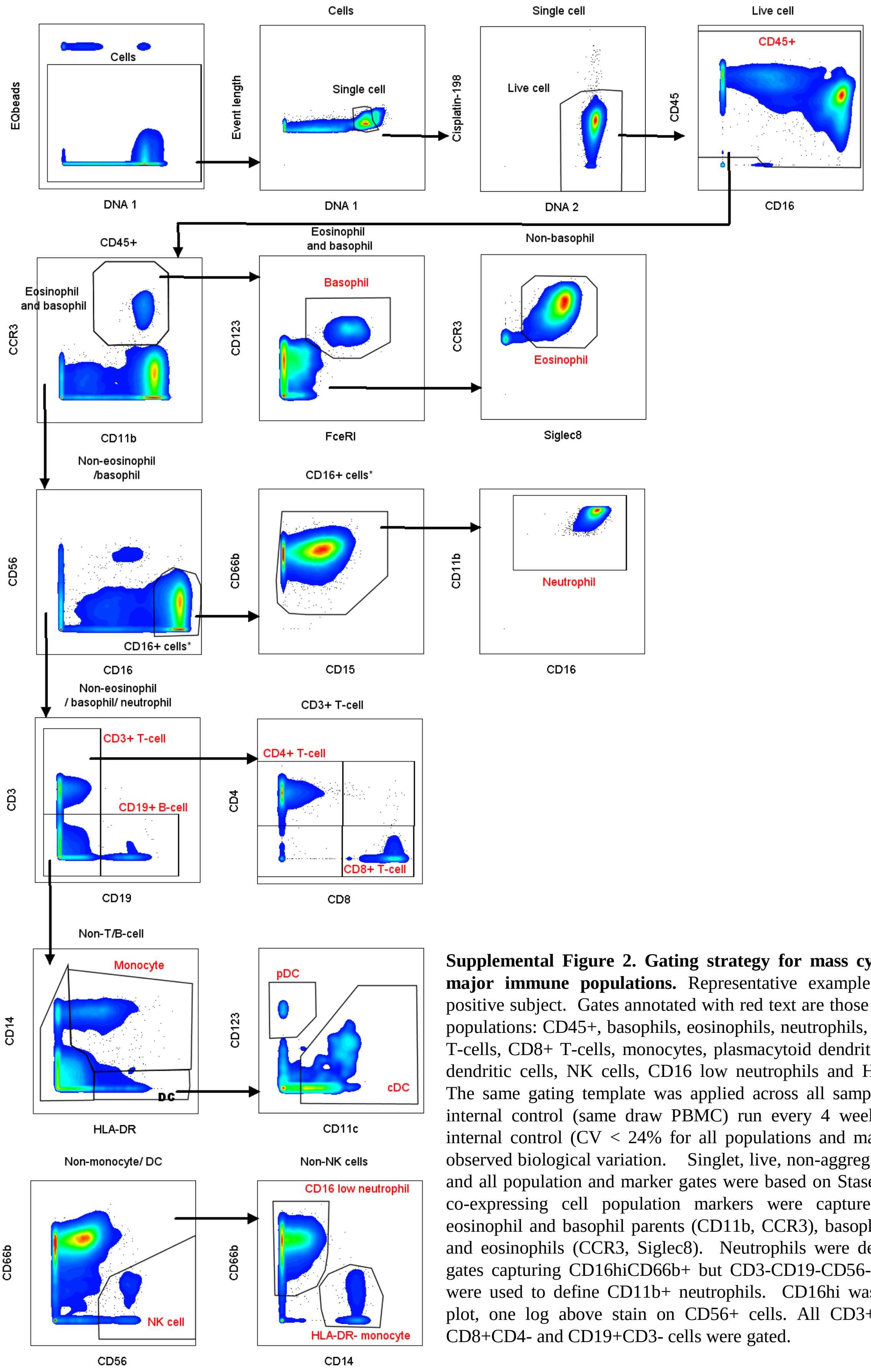

**Supplemental Figure 2. Gating strategy for mass cytometry analysis of major immune populations.** Representative example from a COVID-19 positive subject. Gates annotated with red text are those defining the reported populations: CD45+, basophils, eosinophils, neutrophils, CD3+ T-cells, CD4+ T-cells, CD8+ T-cells, monocytes, plasmacytoid dendritic cells, conventional dendritic cells, NK cells, CD16 low neutrophils and HLA-DR- monocytes. The same gating template was applied across all samples in addition to an internal control (same draw PBMC) run every 4 weeks. Variation for the internal control (CV < 24% for all populations and markers) was less than observed biological variation. Singlet, live, non-aggregate gates are standard and all population and marker gates were based on Staser et.al (31). All cells co-expressing cell population markers were captured in the gates for eosinophil and basophil parents (CD11b, CCR3), basophils (CD123, FceR1), and eosinophils (CCR3, Siglec8). Neutrophils were defined using Boolean gates capturing CD16hiCD66b+ but CD3-CD19-CD56- and HLA-DR- cells were used to define CD11b+ neutrophils. CD16hi was defined by contour plot, one log above stain on CD56+ cells. All CD3+CD19-, CD4+CD8-, CD8+CD4- and CD19+CD3- cells were gated.

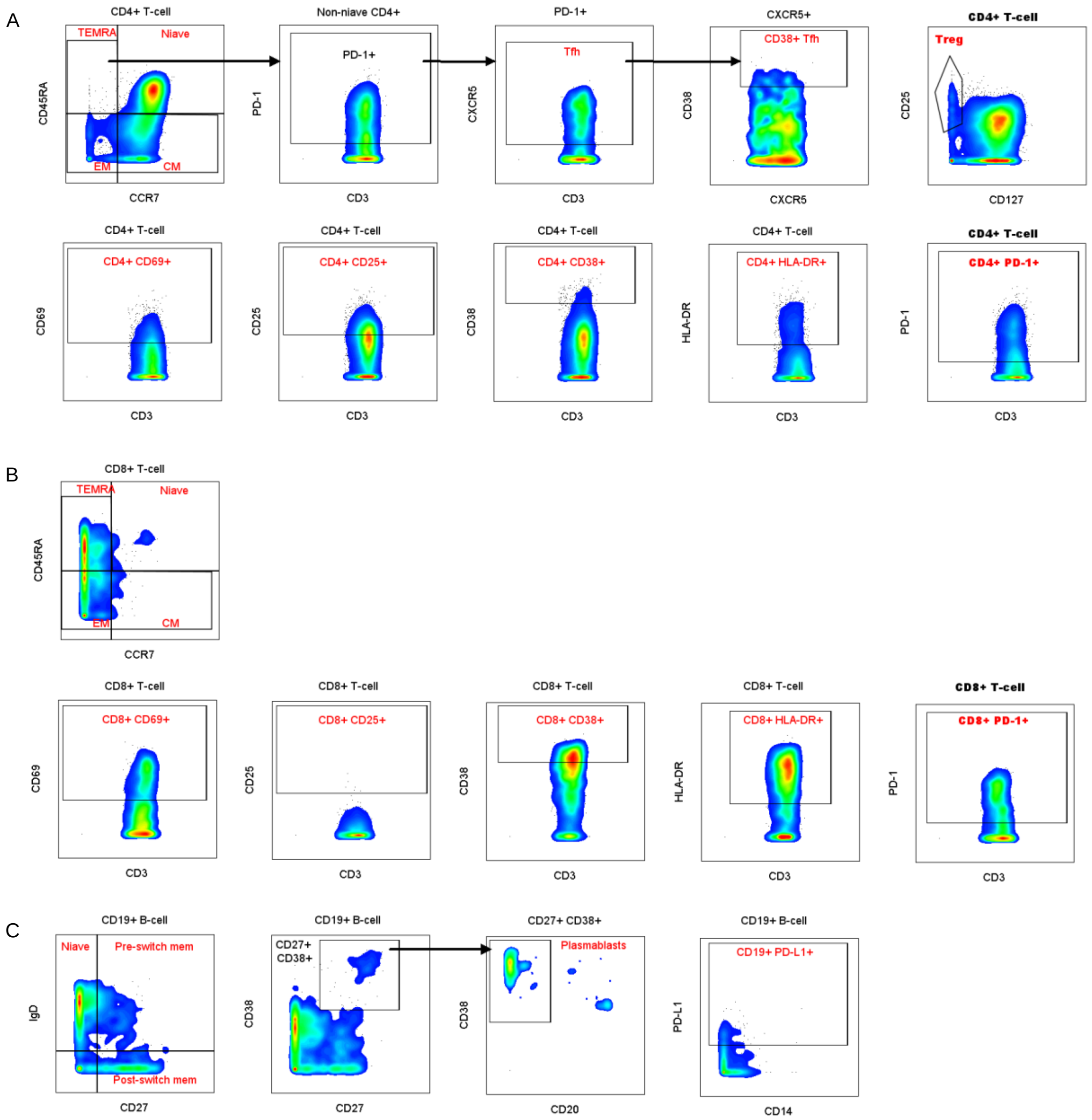

**Supplemental Figure 3. Gating strategy for mass cytometry analysis of T-cell and B-cell subsets.** Representative example from a COVID-19 positive subject. Gates annotated with red text are those defining the reported populations. (**A**) Characterization of CD4+ T-cell subsets and activation. (**B**) Characterization of CD8+ T-cell subsets and activation. (**C**) Characterization of B-cell subsets and activation. The same gating template was applied across all samples based on Staser et.al (31). CCR7xCD45RA quadrant gates for CD3+ CD4 T cells were set based on a tight gate on the naive CCR7+CD45RA+ population. This gate was applied to CD8 T cells. PD-1, CXCR5, CD38, HLA-DR, CD69, and CD25 gates were based on naive cells of controls that lacked expression of these markers and were never adjusted when applying to study samples. CD27xIgD quadrant gates for CD19+ B cells were set based on the naive IgD+CD27- population. PDL-1 was based on a negative control and was never adjusted when applying to study samples. To determine gates for activation markers such as CD25, CD69 and CD38 on T cells and PD-L1 on myeloid cells, we first analyzed 12 samples from 6 subjects with moderate COVID-19 and 6 subjects with severe COVID-19. Gates were set based on a comparison between samples that were clearly highly activated and those that were clearly non-activated. These gates were then applied to all samples in the study and used consistently for all populations analyzed. Specifically, gates for CD25, CD38, CD69, HLA-DR, PD-1 and PD-L1 were the same for all cell types where they were applied.

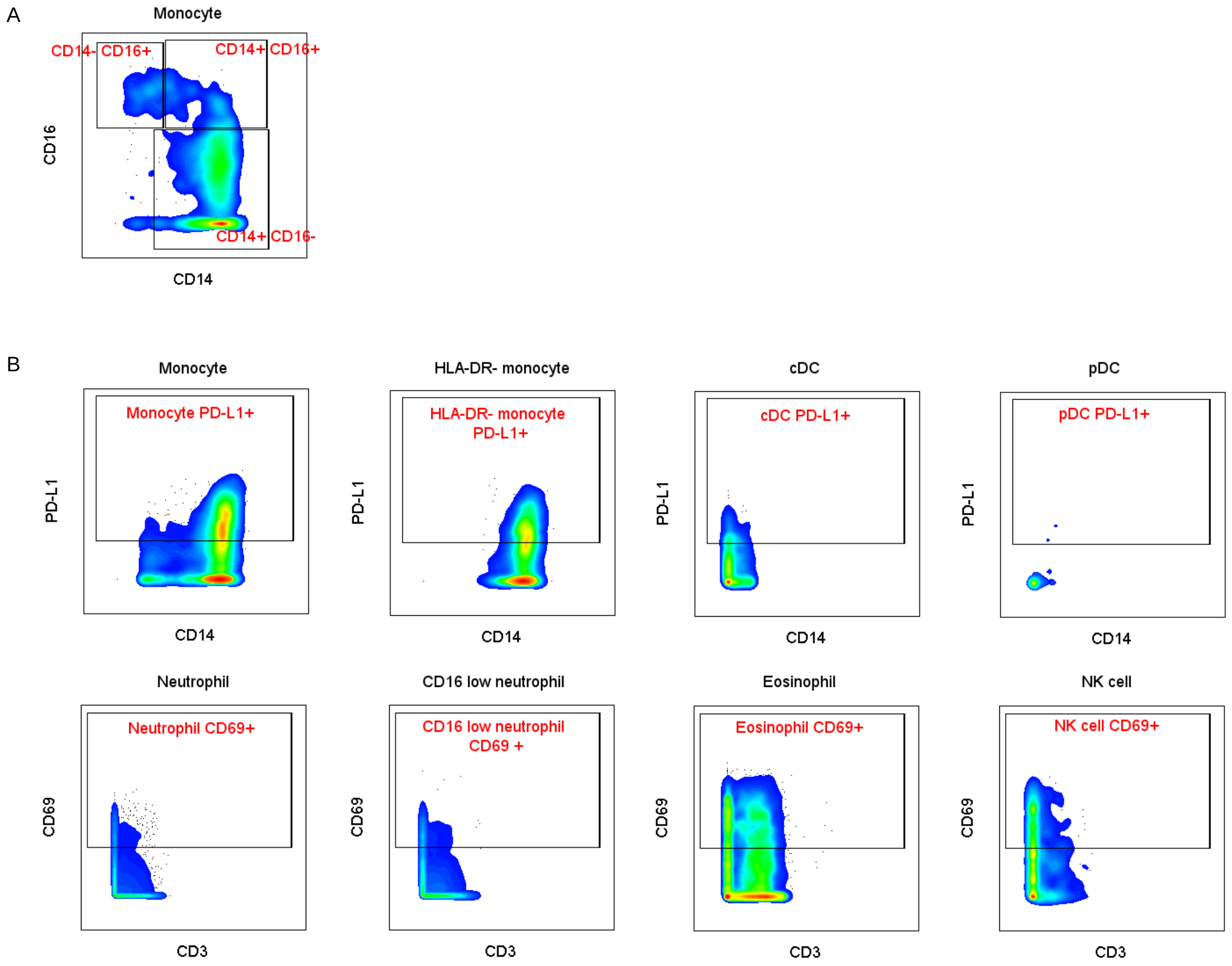

**Supplemental Figure 4. Gating strategy for mass cytometry analysis of monocyte, dendritic cell, neutrophil, eosinophil and NK cell subsets.** Representative example from a COVID-19 positive subject. Gates annotated with red text are those defining the reported populations. (**A**) Characterization of classical (CD14+CD16-), non-classical (CD14loCD16+) and intermediate (CD14+CD16+) monocyte subsets. Gating strategy for these subsets removed cells expressing CCR3, CD15, CD66b, CD3, CD19 and CD56 to define the parent monocyte population. (**B**) Characterization of PD-L1 and CD69 expression by monocytes, dendritic cells, neutrophils, eosinophils and NK cells. Populations were gated as in Supplemental Figures 2 and 3.

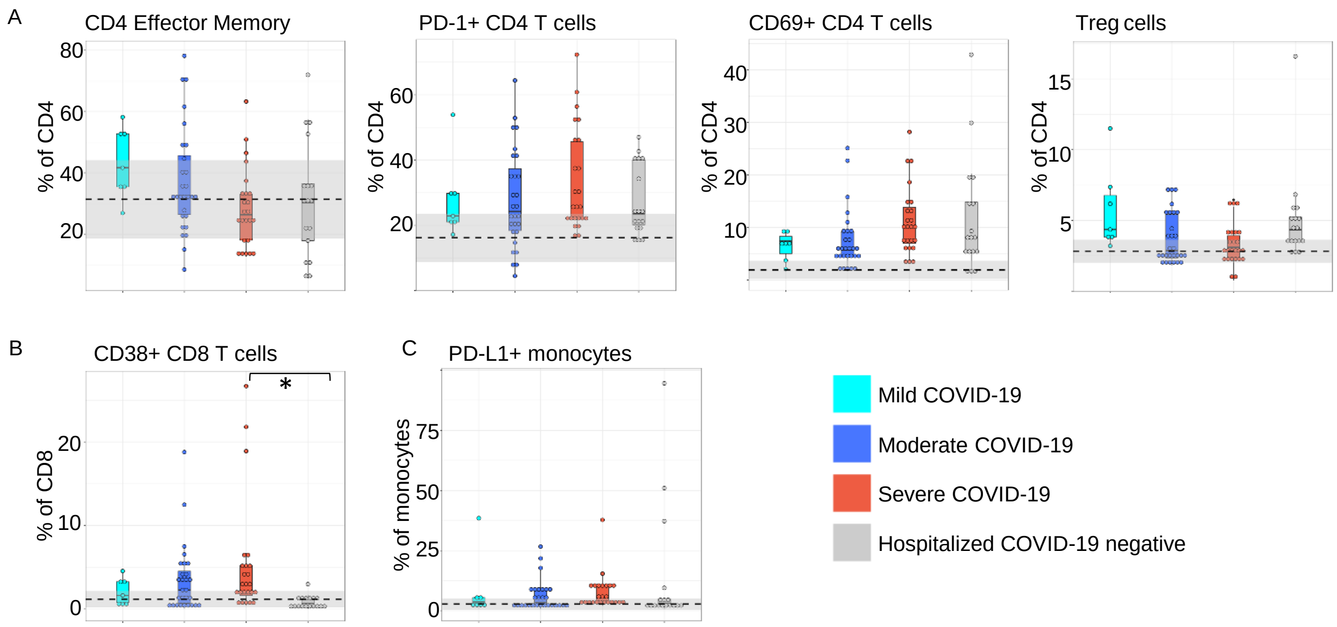

**Supplemental Figure 5. Relative proportions of immune cell sub-types vary by disease severity**. Shown are CyTOF cell frequencies expressed as percentage of parent population. (**A**) CD4 T cell subsets, (**B**) CD8 T cell subset, and (**C**) Monocytes

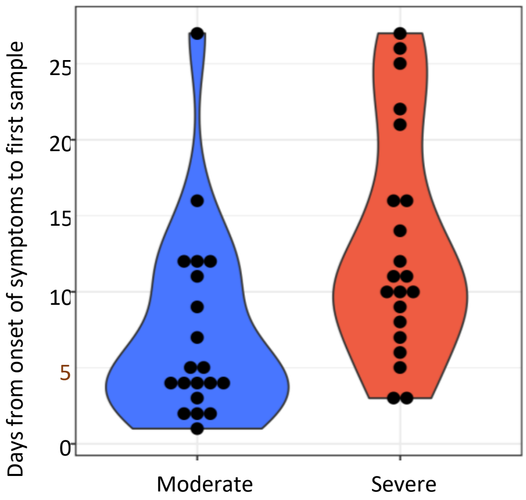

**Supplemental Figure 6.** Sample collection dates relative to the onset of symptoms for moderate and severe COVID-19 subjects with unambiguous onset dates.

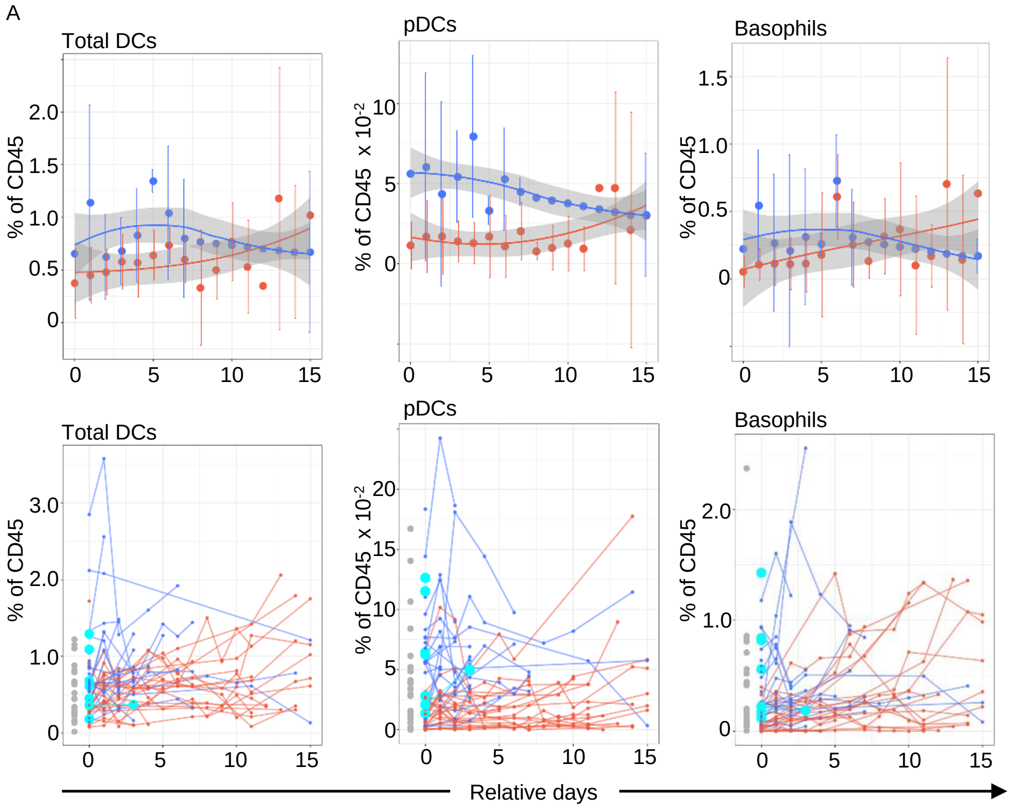

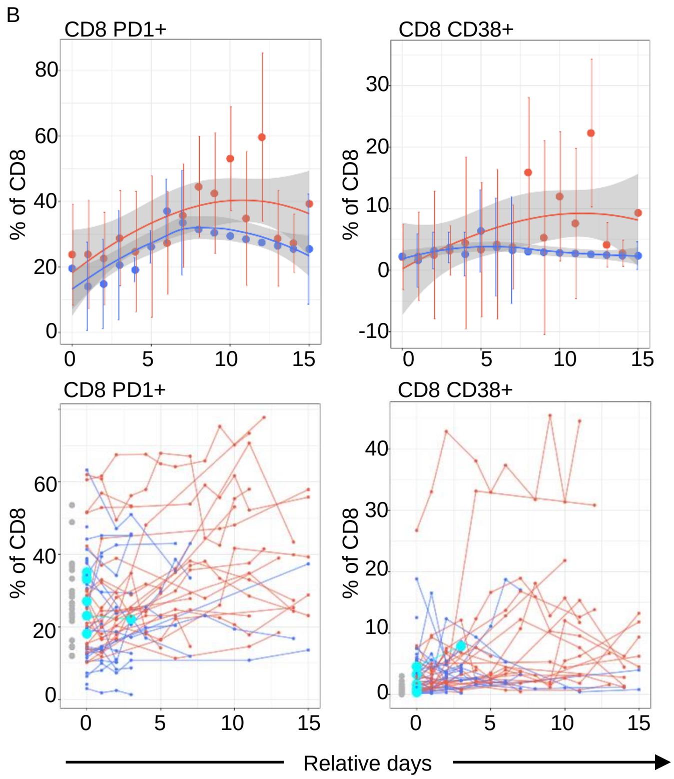

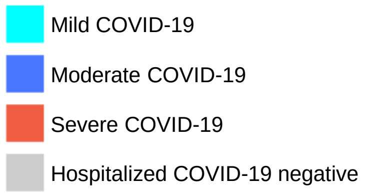

**Supplemental Figure 7**. Smoothed-average, and per-patient longitudinal sampling of immune cell populations measured as percentage of CD45+ cells (**A**) and percentage of CD8+ cells (**B**). The horizontal axes indicate the number of days from first hospitalization. The upper panels in **A** and **B** show smoothed, average trajectories per patient group (see color key). Loess trajectory smoothing was performed on the median values (colored disks) for each group at each time point. Vertical bars indicate +/- 1 standard deviation around the median at each time point. Plot points without vertical error bars are from single data points, or interpolated values used for smoothing. The lower panels show the raw (un-smoothed) trajectories per donor colored by patient group. For comparison, corresponding values for hospitalized COVID-19-negative patients (gray dots) and for patients with mild Covid-19 (cyan dots) are shown on the left-hand side of each panel.

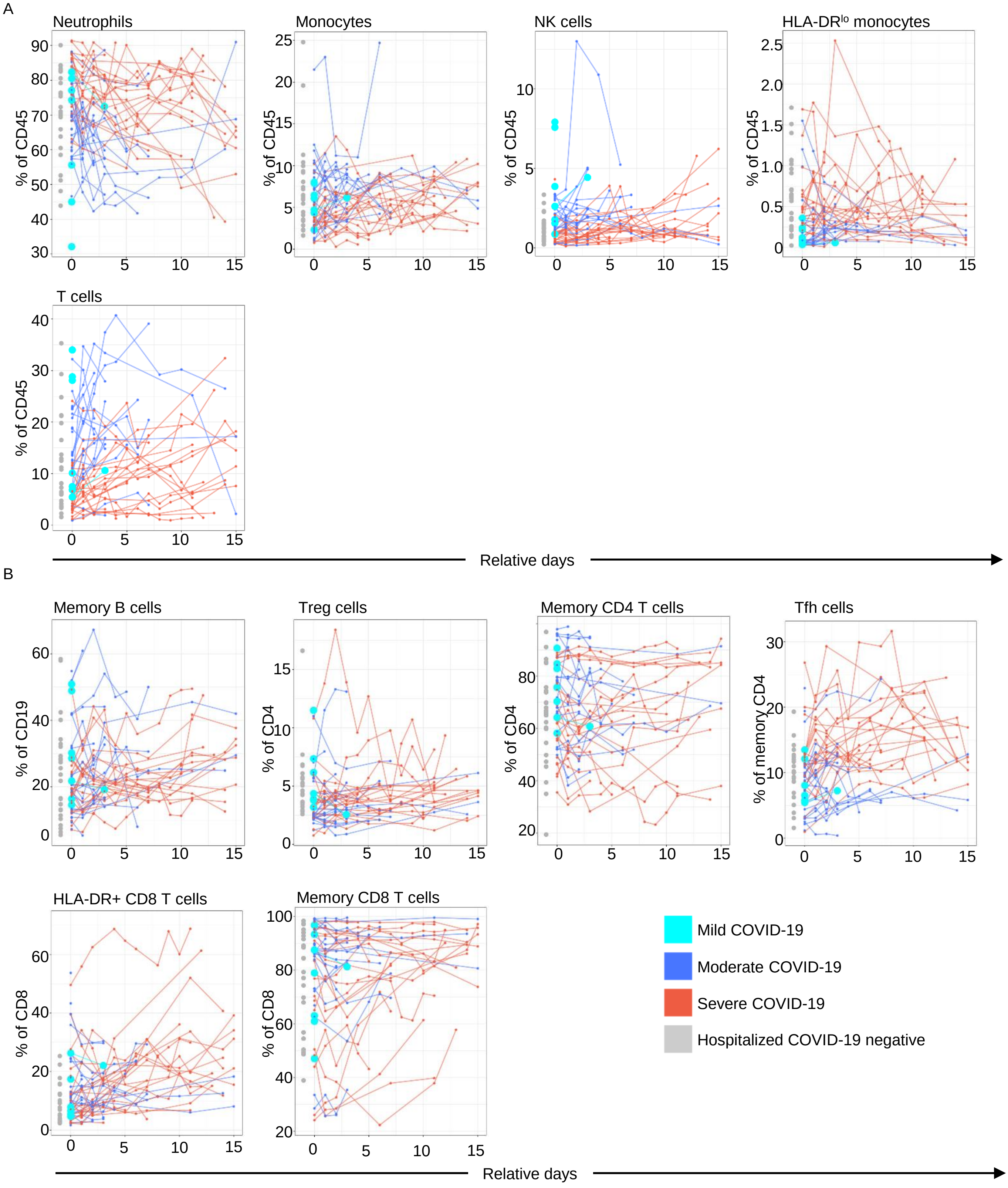

**Supplemental Figure 8**. Longitudinal sampling for individuals and smoothed averages for populations as percentages of CD45 (**A**) and indicated parent populations (**B**). Days (relative) from first hospitalization are shown. Loess trajectory smoothing was performed on the median values (colored disks) for each group at each time point. Vertical bars indicate +/- 1 standard deviation around the median at each time point. Plot points without vertical error bars are from single data points, or interpolated values used for smoothing.

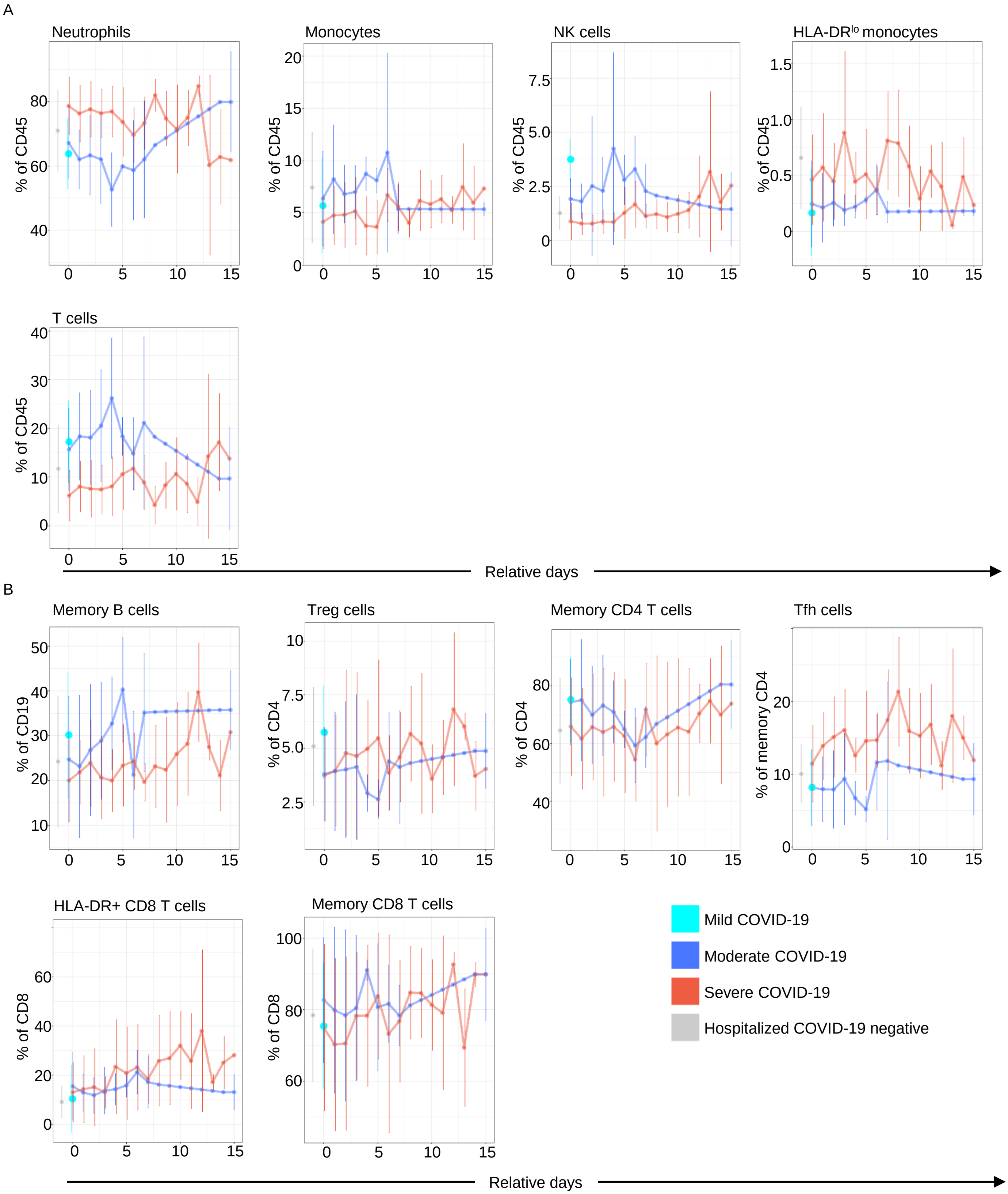

**Supplemental Figure 9**. Un-smoothed time-trajectories of median cell frequencies per patient group for the same cell populations as in Supplemental Figures 8A and 8B.

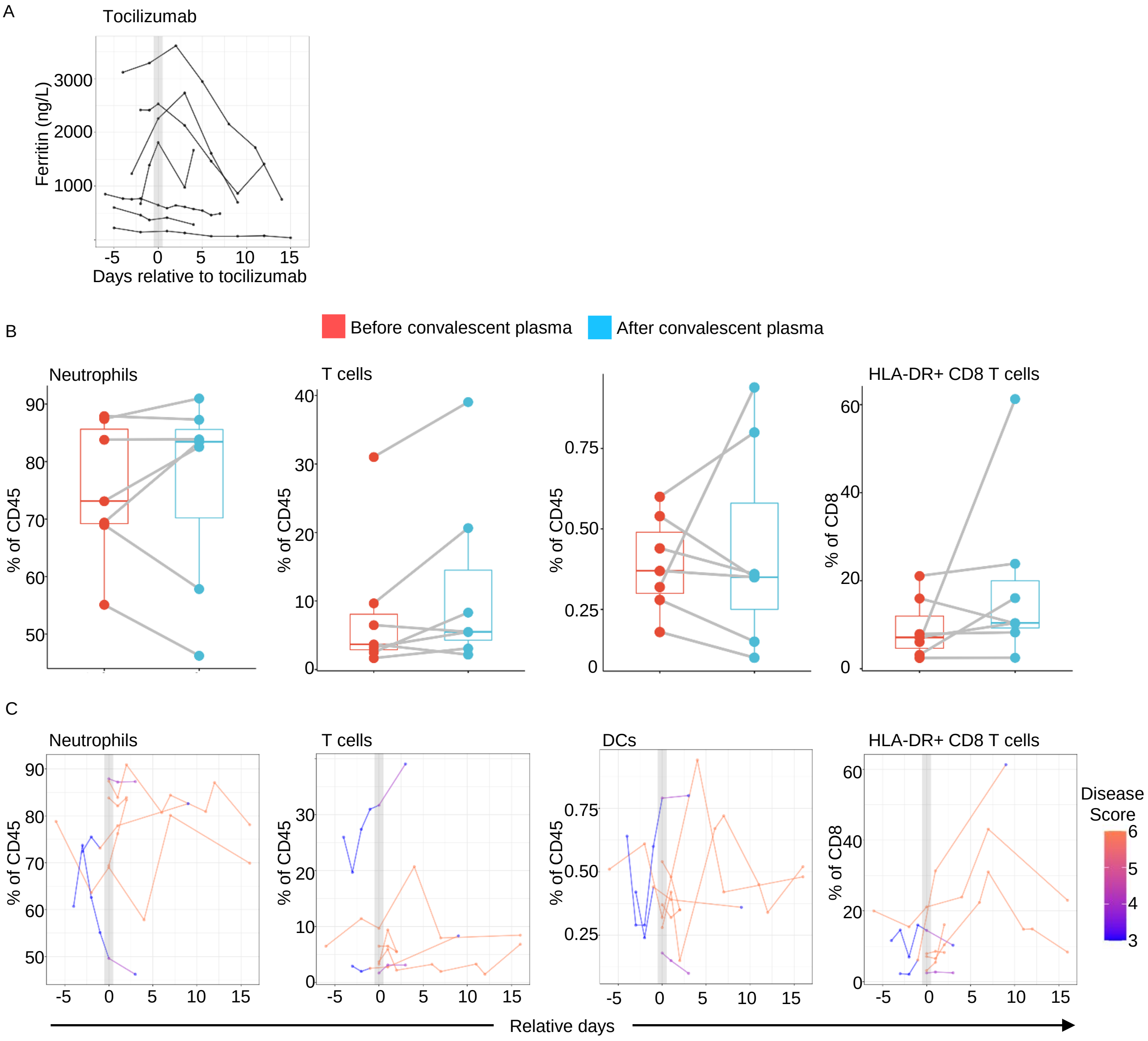

**Supplemental Fig. 10. Analysis of clinical and immune parameters after Tocilizumab and convalescent plasma treatment.** (**A**) Serum Ferritin in patients receiving Tociluzimab. Each line represents an individual patient. (**B**) Plots showing the percent of the indicated populations in convalescent plasma-treated patients before and after treatment, using the same time points used for analysis in Figure 7C see Supplemental Table 3. None of the changes were significant. (**C**) Plots showing all the data points available for convalescent plasma treated patients for the indicated populations shown in (B). Each line represents an individual patient and the color of the line reflects the clinical ordinal score at the time of sampling. Populations were chosen to match those in Figure 10D and 10E.

**Supplemental Table 1. CyTOF panel antibodies**

| Label | Target | Clone | Catalog # | Supplier |
|---|---|---|---|---|
| 89Y | CD45 | HI30 | 3089003B | Fluidigm |
| 141Pr | CD3 | UCHT1 | 3141019B | Fluidigm |
| 142Nd | CD19 | HIB19 | 3142001B | Fluidigm |
| 143Nd | CD123 | 6H6 | 3143014B | Fluidigm |
| 144Nd | CD15 | W6D3 | 3144019B | Fluidigm |
| 145Nd | CD4 | RPA-T4 | 3145001B | Biolegend |
| 146Nd | IgD | IA6-2 | 3146005B | Fluidigm |
| 147Sm | CD11c | Bu15 | 3147008B | Fluidigm |
| 148Nd | FceR1 | AER-37[CRA-1] | 334602 | Biolegend |
| 149Sm | CD127 | A019D5 | 3149011B | Fluidigm |
| 150Nd | CD27 | LG.3A10 | 3150017B | Fluidigm |
| 151Eu | CCR7 | G043H7 | 353202 | Biolegend |
| 152Sm | CD11b | LH2 | 393102 | Biolegend |
| 153Eu | HLA-DR | L243 | 307602 | Biolegend |
| 154Sm | CD38 | HB-7 | 356602 | Biolegend |
| 155Gd | PD-1 | EH12.2H7 | 3155009B | Fluidigm |
| 156Gd | PD-L1 | 29E.2A3 | 3156026B | Fluidigm |
| 158Gd | Siglec 8 | 7C9 | 347102 | Biolegend |
| 159Tb | CCR3 | 5E8 | 310702 | Biolegend |
| 160Gd | CD14 | M5E2 | 3160001B | Fluidigm |
| 161Dy | CD56 | NCAM16.2 | 559043 | BD |
| 162Dy | CD66b | 80H3 | 3162023B | Fluidigm |
| 164Dy | CXCR5 | RF8B2 | 3164029B | Fluidigm |
| 165Ho | CD45RA | HI100 | 304102 | Biolegend |
| 166Er | CD24 | ML5 | 3166007B | Fluidigm |
| 167Er | CD8 | RPA-T8 | 301002 | Biolegend |
| 169Tm | CD25 | 2A3 | 3169003B | Fluidigm |
| 170Er | CD69 | FN50 | 310902 | Biolegend |
| 171Yb | CD141 | M80 | 344102 | Biolegend |
| 172Yb | CD20 | 2H7 | 302302 | Biolegend |
| 175Lu | HLA-A2 | BB7.2 | 343302 | Biolegend |
| 176Yb | HLA-B7 | BB7.1 | 372402 | Biolegend |
| 209Bi | CD16 | 3G8 | 3209002B | Fluidigm |

**Supplemental Table 2. Treatment cohorts demographic and clinical characteristics**

| | Tocilizumab (n=7) | Convalescent plasma (n= 7) |
|---|---|---|
| | **Median (Range)** | **Median (Range)** |
| Age (yrs) | 55 (43-63) | 63 (42-85) |
| Number of Days Hospitalized | 18 (11-40) | 20 (4-60) |
| BMI | 29.8 (26.8-48.9) | 30.4 (18.1-48.9) |
| Disease score at time of experimental treatment | 6 (5-6) | 6 (4-6) |
| Pre-treatment CRP (mg/L) | 285.3 (174.6-350.2) | 215.9 (27.8-323.9) |
| Pre-treatment ferritin (ng/mL) | 1811 (144-3290) | 330 (173-732) |
| Pre-treatment Soluble IL-6 (pg/mL) | 323 (99-1490) | 1032 (51-51512) |
| | **# (%)** | **# (%)** |
| Received care in critical care unit (CCU) | 7 (100%) | 5 (71.4%) |
| Outcome: discharged | 6 (85.7%) | 6 (85.7%) |
| Outcome: deceased | 1 (14.3%) | 1 (14.3%) |
| Female | 5 (71.4%) | 3 (42.9%) |
| **Race** | | |
| Asian | 1 (14.3%) | 0 (0%) |
| White | 1 (14.3%) | 5 (71.4%) |
| Unknown/Other | 5 (71.4%) | 2 (28.6%) |
| **Ethnicity: Hispanic/Latino** | 4 (57.1%) | 2 (28.6%) |
| Exposure to experimental medicine (ever) | | |
| Hydroxychloroquine | 1 (14.3%) | 1 (14.3%) |
| Remdesivir | 7 (100%) | 6 (85.7%) |
| Tocilizumab | 7 (100%) | 2 (28.6%) |
| Convalescent Plasma | 6 (85.7%) | 7 (100%) |

**Supplemental Table 3. Time points analyzed in Figure 10 and Supplemental Figure 10**

| Tocilizumab cohort | Pre-treatment time point (days) | Post-treatment time point (days) |
|---|---|---|
| Patient #1 | -1 | 7 |
| Patient #2 | 0 | 2 |
| Patient #3 | -4 | 5 |
| Patient #4 | -1 | 4 |
| Patient #5 | 0 | 2 |
| Patient #6 | -1 | 2 |
| Patient #7 | -3 | 3 |
| **Convalescent plasma cohort** | | |
| Patient #1 | 0 | 2 |
| Patient #2 | 0 | 4 |
| Patient #3 | -1 | 3 |
| Patient #4 | -1 | 9 |
| Patient #5 | 0 | 3 |
| Patient #6 | 0 | 2 |
| Patient #7 | 0 | 2 |
